# A competition of critics in human decision-making

**DOI:** 10.1101/2020.12.01.407239

**Authors:** Enkhzaya Enkhtaivan, Joel Nishimura, Cheng Ly, Amy Cochran

**Affiliations:** Department of Mathematics, University of Wisconsin, Madison, WI, USA; School of Mathematical and Natural Sciences, Arizona State University, Glendale, AZ, USA; Department of Statistical Sciences and Operations Research, Virginia Commonwealth University, Richmond, VA, USA; Department of Population Health Sciences, University of Wisconsin, Madison, WI, USA

**Keywords:** Decision-Making, Reward Learning, Computational Model, Dopamine, Serotonin, Reaction Time, Risk, Uncertainty

## Abstract

Recent experiments and theories of human decision-making suggest positive and negative errors are processed and encoded differently by serotonin and dopamine, with serotonin possibly serving to oppose dopamine and protect against risky decisions. We introduce a temporal difference (TD) model of human decision-making to account for these features. Our model involves two critics, an optimistic learning system and a pessimistic learning system, whose predictions are integrated in time to control how potential decisions compete to be selected. Our model predicts that human decision-making can be decomposed along two dimensions: the degree to which the individual is sensitive to (1) risk and (2) uncertainty. In addition, we demonstrate that the model can learn about reward expectations and uncertainty, and provide information about reaction time despite not modeling these variables directly. Lastly, we simulate a recent experiment to show how updates of the two learning systems could relate to dopamine and serotonin transients, thereby providing a mathematical formalism to serotonin’s hypothesized role as an opponent to dopamine. This new model should be useful for future experiments on human decision-making.

## Introduction

Temporal difference (TD) learning has enjoyed tremendous support as a conceptual framework for understanding how people make decisions and what might be computed in the brain. TD learning is also supported by studies suggesting that prediction errors derived from a TD model are encoded in dopamine transients [1–5]. Recent theories and experiments, however, suggest that TD models can oversimplify human decision-making in meaningful ways [6–9]. In particular, models that are sensitive to risk or track multiple errors are better able to predict what decisions a person selects [10–17], yet the brain structures involved are not completely known. Similarly, a single-neurotransmitter based circuit, where positive concentrations match prediction-error, would struggle to encode large negative updates [17], and indeed, recent evidence suggests that serotonin may play a complementary role [6,7,18–22]. Our goal was to develop and analyze a simple computational model that resolves and unites these observations. Our proposed model involves dual critics, composed of an optimistic dopamine-like TD learner and a pessimistic serotonin-like TD learner, who compete in time to determine decisions.

TD learning was designed to utilize simple mathematical updates to produce a system that learns how to make decisions [23]. Such models decompose decision-making into two processes: a learning process, which updates how one values a decision, and a decision process, which selects decisions according to how they are valued. These models, including the model of Rescorla and Wagner [24], can learn about reward expectations through updates that are linear in a single prediction error, but are not sensitive to risk or track multidimensional errors.

One reason to expect risk-sensitivity is there is asymmetry in how negative versus positive errors are updated. Dopamine transients, for example, have been found to respond more greatly to positive prediction errors than negative prediction errors [25]. From a biological perspective, this is not surprising. Dopamine neurons have low baseline activity, which imposes a physical limit on how much their firing rates can decrease because firing rates are non-negative [26]. This limit suggests that dopamine neuron firing rates could not be decreased to encode negative prediction errors to the same degree as they can be increased to encode positive prediction errors. If this is true, then the outsized influence of positive prediction errors would inflate the valuation of decisions — colloquially referred to as “wearing rose-colored glasses.”

Computational models capture risk-sensitivity by weighing positive prediction errors differently than negative prediction errors, usually accomplished with separate learning rates for positive and negative prediction errors. These models are referred to as *risk-sensitive*, because they result in decision-making that is sensitive to large gains (i.e. *risk-seeking*) or large losses (i.e, *risk-averse*). Taken to an extreme, risk-seeking involves pursuing best possible outcomes, whereas risk-aversion involves avoiding worse possible outcomes [27]. For comparison, traditional TD learning is considered *risk-neutral* because it focuses on maximizing average (long-term discounted) rewards, so that all rewards, regardless of size, are weighted equally. Risk-sensitive models are frequently found to fit data better than risk-neutral models [15–17]. Importantly, differences in risk-sensitivity, substantiated by a risk-sensitive learning model, is thought to underlie certain differences between individuals with and without psychiatric disorders [28, 29].

The multidimensional aspect of TD-based human decision-making is supported by recent studies. One theory is that serotonin also encodes prediction errors but acts as an opponent to dopamine [6, 7]. In Moran et al, for example, serotonin transients were found to respond to prediction errors in an opposite direction of dopamine transients [7]. Their results were consistent with the hypothesis that serotonin protects against losses during decision-making [7] or more broadly, plays a role in avoidance behavior [21, 30, 31]. Furthermore, a recent study even suggests dopamine is capable of capturing a distribution of prediction errors, the computational benefit of which is that the reward distribution can be learned rather than just its average and variance [9]. Other conceptual frameworks suggest individuals keep track of multiple prediction errors as a way to capture reward uncertainty in addition to expected rewards [10–14].

In this paper, we introduce and analyze a new model of human decision-making, which we call the Competing-Critics model, which uses asymmetrical and multidimensional prediction errors. Based on a TD learning framework, the model decomposes decision-making into learning and decision processes. The learning process involves two competing critics, one optimistic and another pessimistic. The decision process integrates predictions from each system in time as decisions compete for selection. In what follows, we explore through simulation whether our model can capture ranges of risk-sensitive behavior from risk-averse to risk-seeking and can reflect reward mean and variance. Further, we use this model to make predictions about reaction times and about uncertainty-sensitivity in terms of the degree to which reward uncertainty influences a person’s consideration of multiple decisions. Lastly, we show how prediction errors in the Competing-Critics model might relate to dopamine and serotonin transients in the experiments of Kishida *et al* [8] and Moran *et al* [7]. Considering the simplicity of this model and its ability to synthesize several theories and experimental findings, this model should be useful as a framework for future human decision-making experiments, with potential to provide both predictive power and mechanistic insight.

## Modeling

We introduce a model of human decision-making that relies on two competing learning systems. Figure 1 provides a high-level view of the proposed model in a simple example in which an individual makes decisions between two choices. Here the individual learns to value their decision by weighing prior outcomes observed upon selecting each choice, denoted by *R_t_*, in two different systems. The first learning system weighs better outcomes more heavily than worse outcomes, which effectively leads to a more optimistic valuation of outcomes, denoted by *Q*^+^. The second learning system does the opposite: weighs worse outcomes more heavily than better outcomes, leading to a more pessimistic valuation of outcomes, denoted by *Q*^−^. We remark that both values, *Q*^+^ and *Q*^−^, are assumed to be updated according to prediction errors 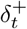 and 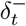, following common risk-sensitive temporal difference (TD) learning frameworks described below.

**Fig 1.**
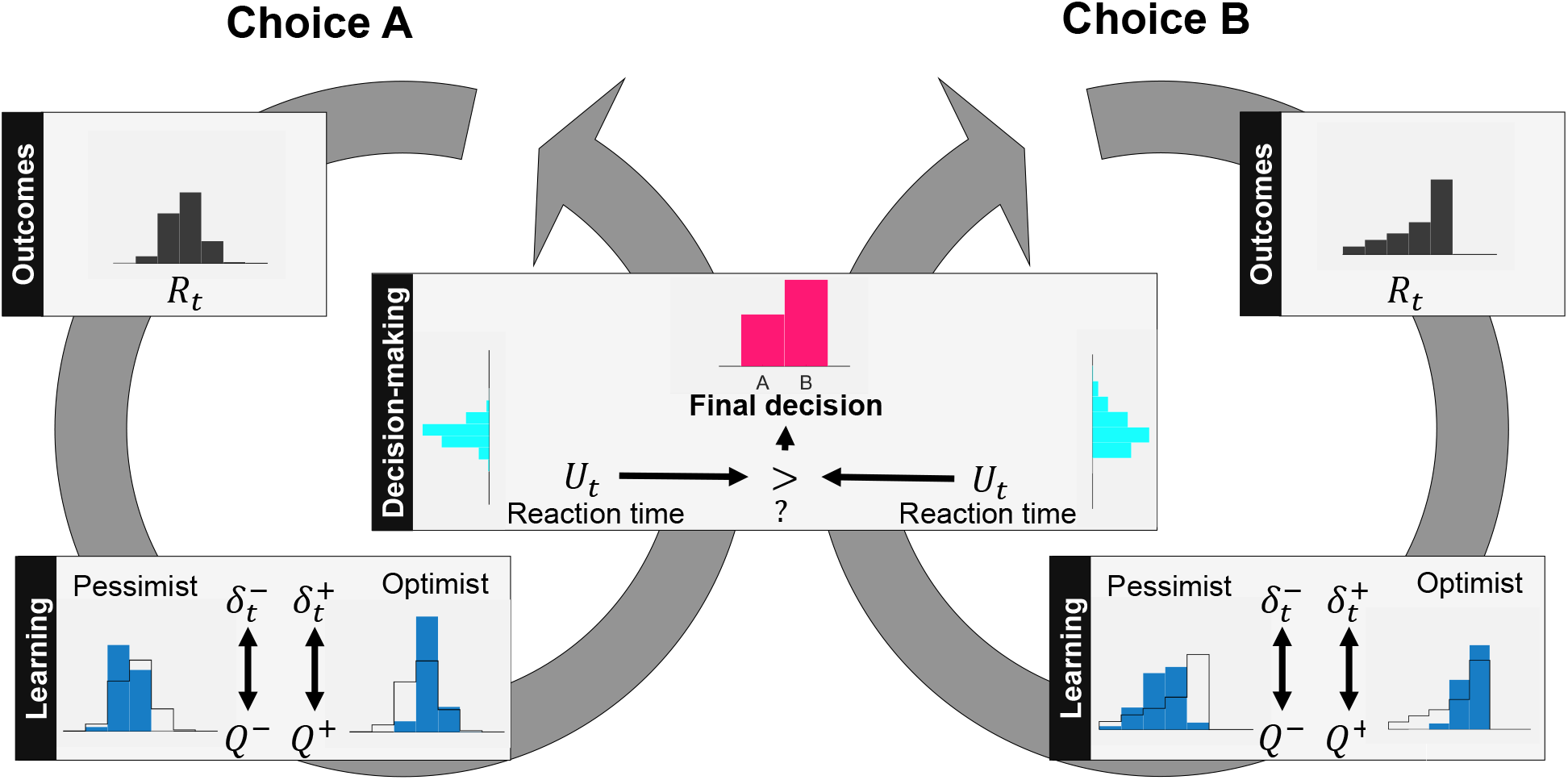
High-level view of proposed model in an example with two choices. For each choice, the distribution of rewards *R_t_* (gray histograms) is learned by competing critics through the updates 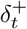 and 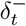. One system is optimistic, upweighting large rewards, and another is pessimistic, downweighting large rewards (blue histograms). As a result, each choice is associated with multiple values *Q*^−^ and *Q*^+^. To determine which choice is selected, a random variable *U_t_* is drawn for each choice uniformly from (*Q*^−^, *Q*^+^) (teal histograms). The largest *U_t_* determines which choice is selected and when the decision is made.

An individual who relied solely on the first learning system to make decisions would be considered *risk-seeking* due to the outsized influence of better outcomes. Similarly, an individual who relied solely on the second system to guide decisions would be considered *risk-averse* due to the outsized influence of worse outcomes. Our model, however, supposes both of these competing learning systems contribute to decision-making in the following way. For each choice, the risk-seeking learning system sends a *go* signal to the individual to signify that this choice is viable, with larger *Q*^+^ values corresponding to earlier signals. Afterwards, the risk-sensitive learning system sends a *no-go* signal to the individual to signify that this choice is no longer viable, with small *Q*^−^ associated with later signals. For simplicity, the individual is assumed to select the respective choice at any time between these two signals, provided no other choice has been selected or choice exploration has been pursued. Hence, both go and no-go signals determines how likely each choice is selected. For example, choices whose go signal is initiated after a no-go signal of another choice will never be selected except for exploration. Put differently, any choice when valued optimistically is still worse than another choice valued pessimistically will not be selected except for exploration. We now proceed to formalize this conceptual framework.

### Setting

Our model will describe psychological experiments that have the following decision-making scenario. The scenario starts at the initial state *S*_0_ on which the participant bases their action *A*_0_, which brings in a numerical reward *R*_1_. Consequently, the participant finds itself in the next state *S*_1_ and selects another action *A*_1_, which brings in a numerical reward *R*_2_ and state *S*_2_. This process then repeats until the participant makes *T* decisions, yielding a sequence of observations collected for each participant of the form:

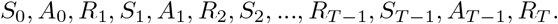

Above, observations fall into three types on a given trial *t*: the *state* that the participant visits, denoted by *S_t_*, the *action* that the participant takes when visiting state *S_t_*, denoted by *A_t_*, and the subsequent *reward*, *R*_*t*+1_, that a participant receives upon visiting state *S_t_* and taking action *A_t_*. For simplicity, let us assume that both the space of possible states 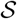 and the space of possible actions 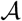 are discrete. Further, assume the experiment defines subsequent rewards and states as a function of the current state and action according to a Markov transition probability

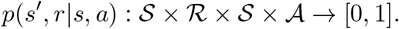

An experiment described above constitutes a (discrete-time, discrete-state) Markov Decision Process (MDP).

Several popular experiments can be described by the setting above. The Iowa Gambling Task (IGT), for example, asks participants to repeatedly chose between four decks labelled A to D. After each choice, they gain and/or lose monetary rewards (Fig 2). Hundreds of studies have used the IGT to evaluate human decision-making [32]. Initial findings found healthy controls would learn to select “good” decks (Decks C and D), so-called because, on average, they yielded a net gain [33]. By contrast, individuals with a damaged prefrontal cortex would continue to select “bad” decks (Decks A and B) despite yielding net losses on average. Selecting bad decks is considered a marker of impaired decision-making.

**Fig 2.**
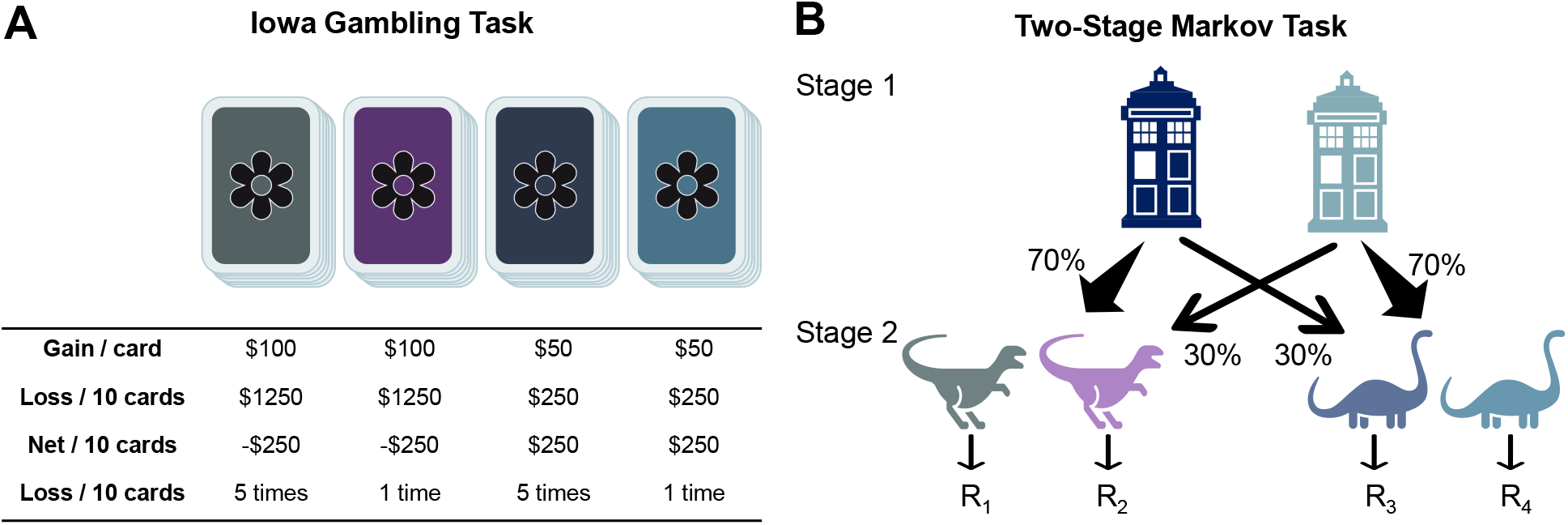
Popular tasks for evaluating decision-making: (**A**) Iowa Gambling and (**B**) Two-Stage Markov tasks. Both tasks can be formulated as Markov decision processes.

The above formulation can be recovered for the IGT with *A_t_* ∈ {*A, B, C, D*} capturing the selected desks, *S_t_* ∈ {1} capturing a trivial case with only one state, and *R_t_* capturing the summed gain and loss per trial. In particular, we will simulate *R_t_* as independent draws from a distribution that depends on the selected deck and matches characteristics in Fig 2**A** on average. For example, *R_t_* is drawn uniformly from {$50, $0} when Deck C is selected. We point out that the IGT is a specific case of a multi-armed bandit, which is a common framework to evaluate human decision-making and can also be described by the above setting.

Another example is the two-stage Markov task [34], in which a participant repeatedly selects images over two stages. Participants are presented one of three pairs of images at a given stage. At the first stage, all participants are shown the first pair of images and have the option to choose either the left or right image. After choosing an image, participants are shown the second or third pair of images, with the pair selected randomly according to probabilities that depends on their first stage selection. Participants then select an image on the second stage and receive monetary rewards. This task is used in experiments to determine the degree to which individuals are learning about the common (*p* = 0.7) versus rare (*p* = 0.3) transition associated with each action in stage 1. To mark this type of learning, the authors point to the probability of staying on the same first-stage decision depending on the type of transition (common vs. rare) and whether or not the person was rewarded on the second stage. In particular, if a person was learning about the transitions and was rewarded, then they were expected to see a drop in the stay probability when a common versus rare transition occurred. If if a person was learning about the transitions and was not rewarded, then they were expected to see a jump in the stay probability when a common versus rare transition occurred.

In terms of the above formulation, this task gives rise to actions *A_t_* ∈ {left, right} representing selected images, states *S_t_* ∈ {1, 2, 3} capturing presented image pairs, and rewards *R_t_* capturing rewards after image selection with rewards after the first stage set to zero. Despite the multi-stage aspect of this task, here *t* counts the total number of actions so that *t* = 0 refers to the first time that a participant takes an action in the first stage and *t* = 1 refers to the first time that a participant takes an action in the second stage.

### Temporal difference (TD) learning

In the setting described above, human decision-making is often modeled using TD learning. One widely-known algorithm for TD learning is called SARSA, so-named for its explicit use of the sequence of observations {*S*_0_, *A*_0_, *R*_1_, *S*_1_, *A*_1_,…}. This algorithm supposes that the agent, i.e., the participant in a psychological experiment, tries to learn the “value” of their actions as a function of a given state in terms of future rewards. This notion gives rise to a state-action value function *Q*(*s, a*) mapping states 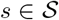 and actions 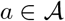 to a real number that reflects the value of this state-action pair. A SARSA learner updates this state-action function according to their experiences:

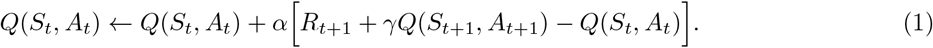

Here, the learner has just taken action *A_t_* in state *S_t_*, receiving the immediate reward *R*_*t*+1_ and transitioning to a new state *S*_*t*+1_, where upon they take action *A*_*t*+1_. A learning rate *α* accounts for the extent to which the new information, i.e. their reward and the new state-action value, overrides old information about their state-action value function. For instance, one can see that if *α* = 0, there is no overriding - the estimate stays the same. The discount parameter *γ* weighs the impact of future rewards. A discount parameter *γ* = 0 would mean the learner does not care about the future at all, while *γ* = 1 would mean the learner cares about the sum total of future rewards (which may even cause the algorithm to diverge).

### Risk-sensitive TD learning

A variant of the SARSA learner allows a learner to be particularly sensitive to smaller, or more negative, rewards, i.e. *risky* situations. In particular, a risk-sensitive SARSA learner weighs the prediction error, which is given by

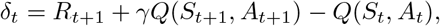

differently depending on whether the prediction error is positive or negative. This yields the following update:

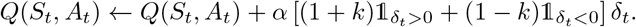

The parameter *k* controls the degree to which the learner is risk sensitive. If *k* = 0, then the learner weighs positive and negative prediction errors equally, in which the updates are the same as before and we say the learner is *risk-neutral*. If *k* < 0, then negative prediction errors are weighed more than positive prediction errors. In this case, smaller rewards have a stronger influence relative than larger rewards on the state-action value function *Q*, resulting in a learner who is considered *risk-averse*. Similarly if *k* > 0, the reverse is true: larger rewards have a stronger influence relative to smaller rewards, and the learner is considered *risk-seeking*.

### A learning model with competing critics

With the introduction of risk-sensitive TD learning, we can consider a range of learning behaviors from risk-sensitive to risk-seeking, all modulated by parameter *k* and reflected in the state-action value function *Q*. Researchers are often focused on how pessimism or risk-sensitivity, substantiated by *k*, might vary between individuals. In our model, however, we investigate how risk-sensitivity might vary within individuals. Specifically, we consider two learning systems, one pessimistic (risk-adverse) and one optimistic (risk-seeking).

Our model captures two competing critics by keeping track of two state-action value functions, *Q*^+^ and *Q*^−^, and updated each function according to:

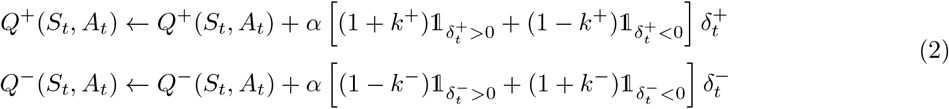

with prediction errors given by

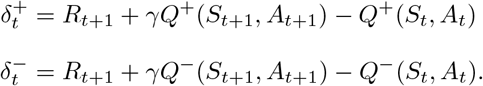

For simplicity, we initialize *Q*^+^ and *Q* to zero. Parameters *k*^+^, *k*^−^ are assumed to lie in [0, 1]. Large *k*^+^ controls the degree to which the learner is risk-seeking and *k*^−^ controls the degree to which the learner is risk-sensitive. It is important to point out that we are also not the first to consider multiple risk-sensitive TD learning systems. This idea was recently put forth in [9], where multiple risk-sensitive TD learning systems were thought to be encoded in multiple dopamine neurons. We are also not the first to consider dual competing systems [6, 18, 35, 36]. In the opposing actor learning model in [35], for example, prediction error from a single learning system controls the dynamics of G (“go”) and N (“no-go”) systems, which in term are combined linearly to determine decisions. Since it may not be obvious why this model differs from our proposed model, we discuss in the Supplement how the update equations of the two models differ in important ways, resulting in significantly different behaviors and predictions.

### A decision-making model with competing critics

Now that we have a model of learning, namely *Q*^+^ and *Q*^−^, it is sensible to consider how the agents makes decisions based on what they have just learned. This means that the individual has to make the decision of choosing from the available actions, having obtained pessimistic and optimistic estimates for action-value pairs.

A naive approach is what is called the greedy method, meaning that the action with the highest value is chosen. This approach, however, does not account for actions with multiple values (e.g., optimistic and pessimistic values) nor does it allow the individual to do any exploration, during which they might discover a more optimal strategy. A way to incorporate exploration into decision-making is to act greedy 1 – *ε* of the time and for *ε* of the time, the individual explores non-greedy action with equal probabilities. This method referred to as *ε*-greedy and is used by our model.

To integrate multi-valued actions into a *ε*-greedy method, our model supposes that a random variable *U_t_*(*a*) is selected for each action *a* uniformly from the interval

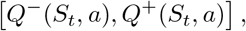

whenever an individual has to make a decision in state *S_t_*. Then whenever the individual acts greedily, they select the action *A_t_* that maximizes *U_t_*(*a*). These decision rules along with learning models comprise Competing-Critics model, which is summarized in Algorithm 1.

#### Algorithm 1: Competing-Critics

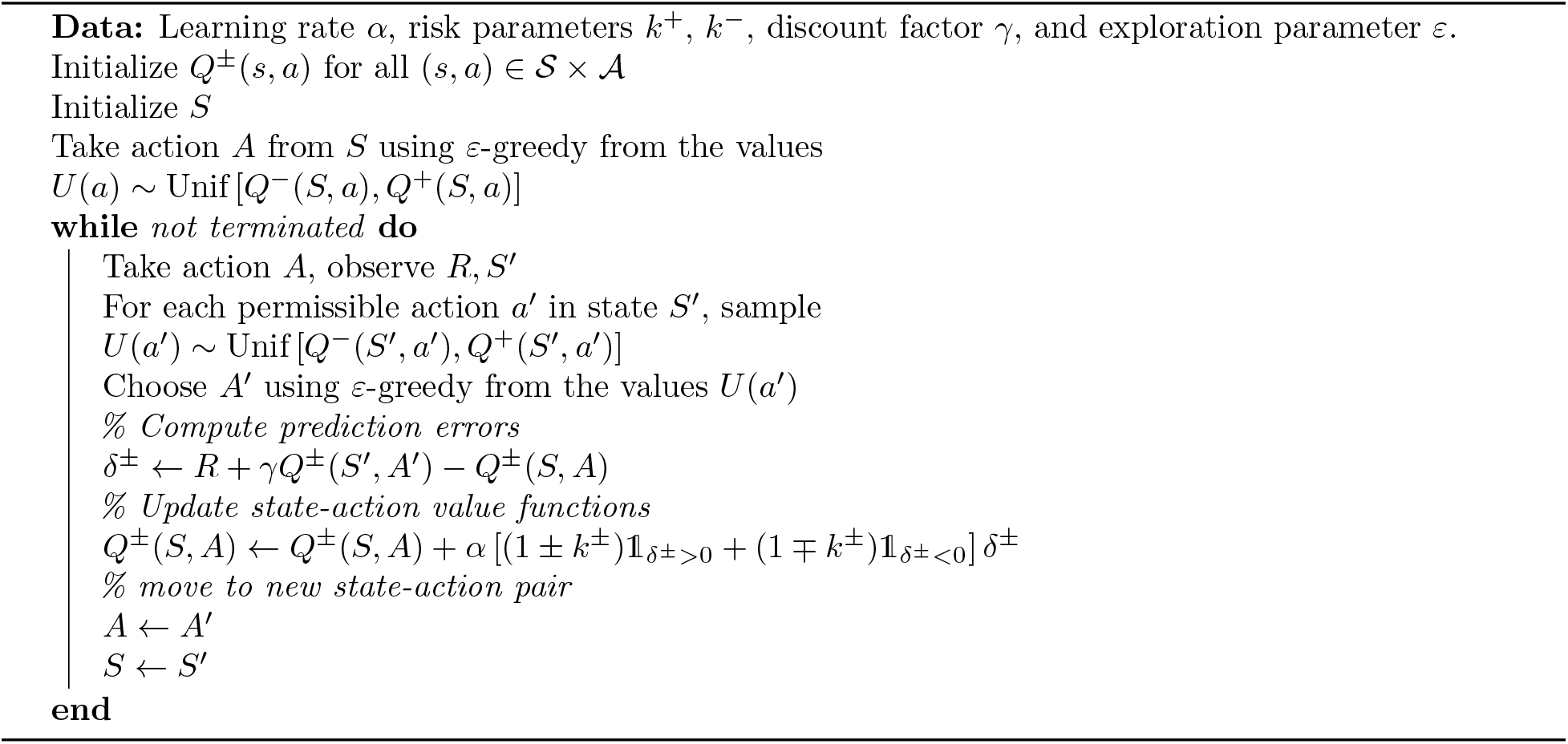

### Simulation experiments

We used simulation to investigate individual behavior in several experiments were they to learn and make decision according to our decision-making model. In particular, we wanted to identify possible vulnerabilities in behavior that arise from a shift in the balance between the internal optimist and pessimist, instantiated by changes in parameters *k*^+^ and *k*^−^. For simplicity, parameters are fixed:

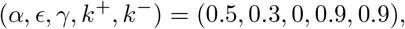

unless otherwise specified. A detailed description of the simulations can be found in the Supplement and at: https://github.com/eza0107/opposite-Systems-for-Decision-Making

### Learning the shape of rewards

Let us first focus on learning behavior by considering the simple case of trivial state and action spaces: 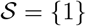 and action 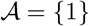. In this case, learning in the Competing-Critics model is determined completely by the distribution of rewards *R_t_*. We considered what an individual would learn given four different Pearson distributions of *R_t_*, with varying mean *μ*, standard deviation *σ*, and skew, while kurtosis was fixed at 2.5. For reference, we also consider the classic *Q* described at Eq. (1).

Figure 3 illustrates what an individual with balanced risk-sensitivity parameters, (*k*^+^, *k*^−^) = (0.9, 0.9), learns over 100 trials. Solid dark lines denote state-action value function averaged over simulations and shaded regions represent associated interquartile ranges (IQRs) for each function. One can immediate notice several things. By design, the optimistic value function *Q*^+^ is on average larger than the neutral value function *Q*, which is larger than the average pessimistic value function *Q*^−^. In addition, the distribution of each value function appears to converge and can capture shifts in mean rewards *μ* and scaling of the standard deviation *σ*. Specifically, the long-term relationship between *Q*^+^, *Q*^−^ and *Q* is preserved when *μ* is shifted from 0.5 to 0.25, whereby all value functions shift down by about 0.25. Further, the gap between *Q*^+^ and *Q*^−^ is halved when *σ* is halved from 0.2 to 0.1; each IQR is also halved. Meanwhile, *Q*^+^ and *Q*^−^ are roughly symmetric around the *Q* when the reward distribution is symmetric (i.e. zero skew), so that the average of *Q*^+^ and *Q*^−^ is approximately *Q*. However, moving skew from 0 to 1 is reflected in both the gap between *Q*^+^ and *Q*, which lengthens, and the gap between *Q*^−^ and *Q*, which shortens.

**Fig 3.**
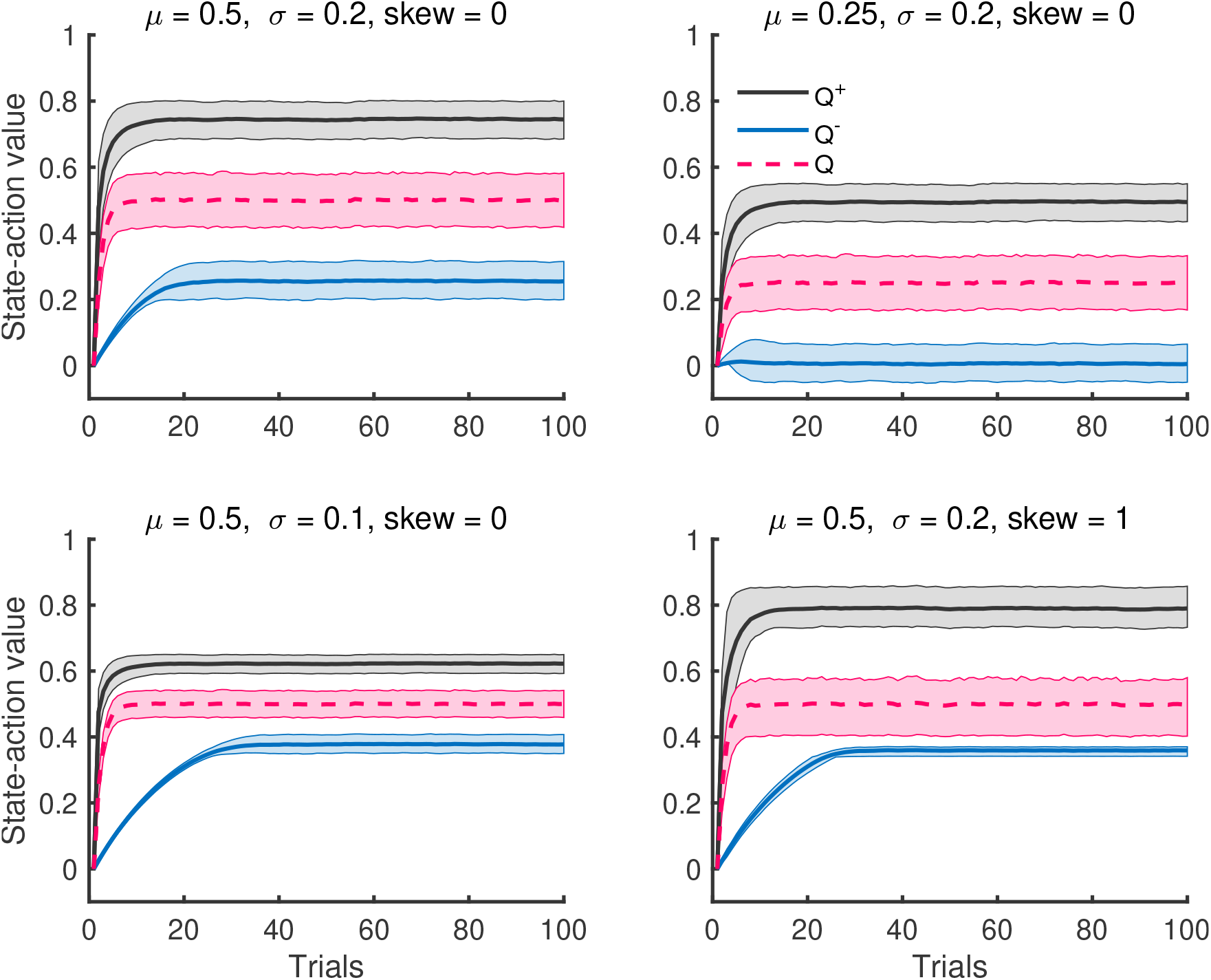
Comparison of mean and interquartile range of state-action value functions over 4000 simulations. The state-action values **Q*^+^* and *Q*^−^ reflect changes in the mean *μ*, standard deviation *σ*, and skew of the reward distribution. Notably, asymptotes of these values shift by 0.25 when *μ* decreases by 0.25, and their gap decreases by 1/2 when *σ* decreases by a factor of 1/2.

Remarkably, the relationship *Q*^+^ > *Q* > *Q*^−^ is also present within a single simulation run (Figure 4). Intuitively, this makes sense because they capture the behaviors of risk-seeking, risk-neutral and risk-sensitive agents, respectively and it turns out that this ordering can be preserved provided *k*^±^ are neither too small or large. See Supplement for the proof of this result. Furthermore, the last subplot also illustrates that introducing a positive skew to the reward distribution *R_t_*, also causes the distribution of *Q*^±^ and *Q* to also have positive skew.

**Fig 4.**
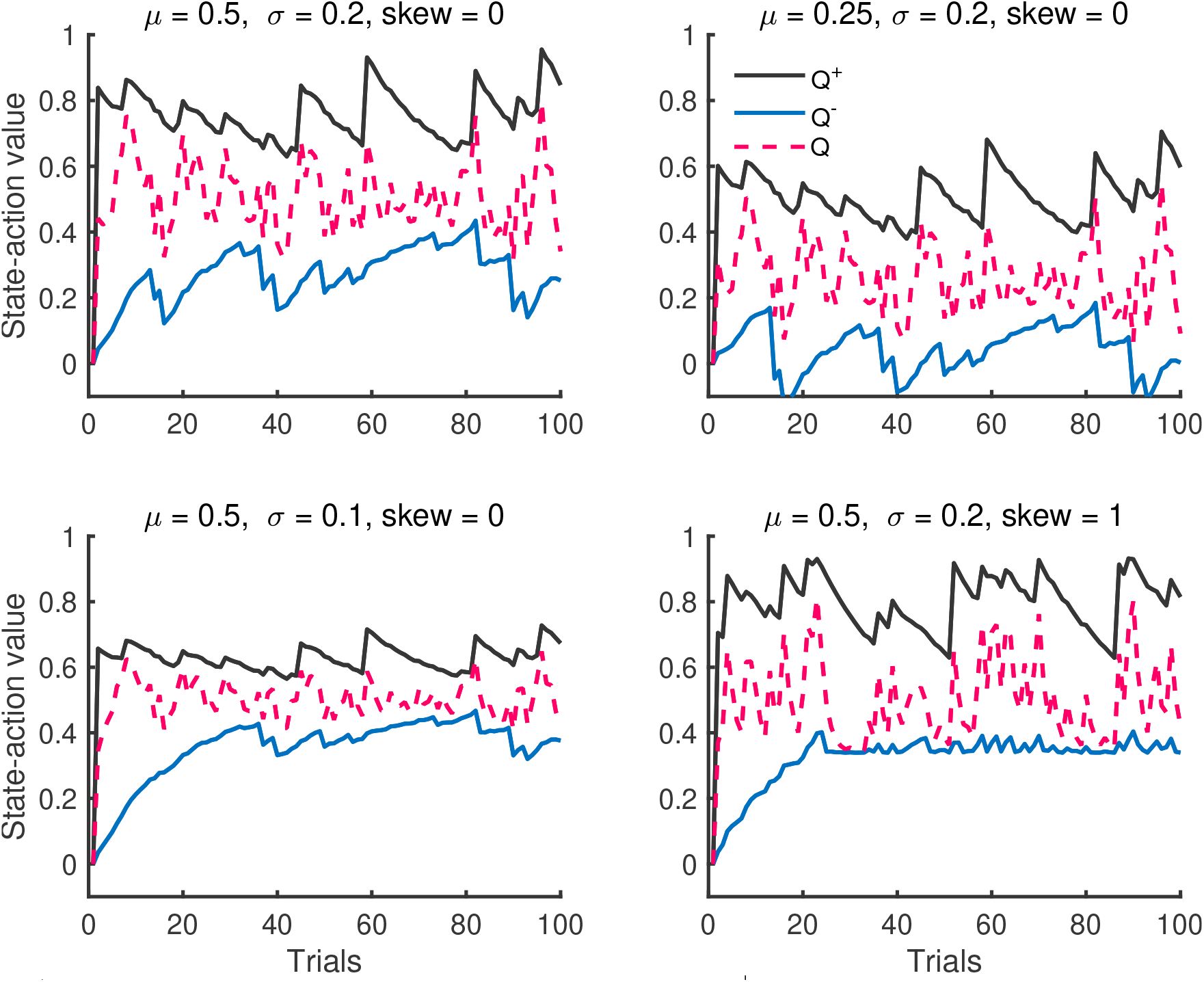
A single simulation run of state-action value functions *Q*^±^ and *Q*. The state-action values preserve the ordering *Q*^−^ < *Q* < *Q*^+^ through the entire run.

Value functions *Q*^+^ and *Q*^−^ are not only modulated with the reward distribution, but also risk-sensitivity parameters. Increasing *k*^+^ moves *Q*^+^ in a positive direction away from the risk-neutral value function *Q*, whereas increasing *k*^−^ moves *Q*^−^ in a negative direction away from the risk-neutral value function *Q*. With *k*^+^ pulling *Q*^+^ in one direction and *k*^−^ pulling *Q*^−^ in the opposite direction, the midpoint of *Q*^+^ and *Q*^−^ is largely influenced by the gap in *k*^−^ and *k*^+^ (Figure 5A). Meanwhile, the gap between *Q*^+^ and *Q*^−^ is largely influenced by the midpoint of *k*^+^ and *k*^−^.

**Fig 5.**
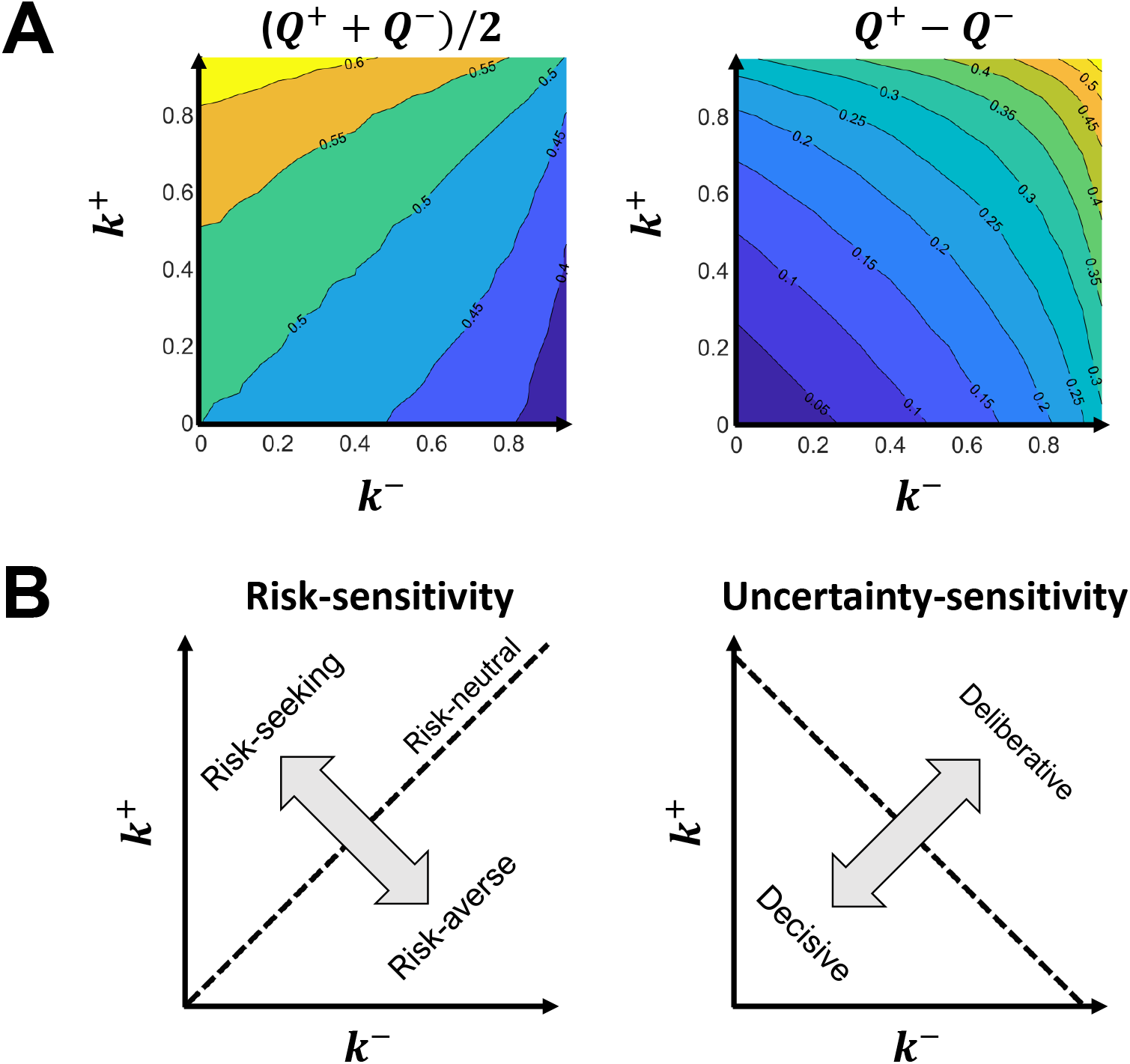
Impact of risk-sensitivity parameters *k*^+^ and *k*^−^ on **A**) the midpoint and gap between *Q*^+^ and *Q*^−^ averaged over 4000 simulations, and **B**) how an individual makes decisions. In particular, the model decomposes decision-making behavior along two axes, a risk-sensitivity and an uncertainty-sensitivity, which are rotated 45° degrees from the *k*^±^ axes. In the simulation, *μ* = 0.5, *σ* = 0.2, and skew = 0.

Since decisions are selected according to the interval between *Q*^+^ and *Q*^−^, the gap in *k*^−^ and *k*^+^ determines the degree to which an individual selects action with high payoffs vs. small losses, i.e. their risk-sensitivity. The midpoint of *k*^−^ and *k*^+^ determines the degree to which reward uncertainty influences whether or not an individual deliberates between actions, i.e. their uncertainty-sensitivity (Figure 5B). This deliberation allows a learner to consider multiple decisions, as one does during exploration but for different reasons. Conceptually, multiple decisions are considered in the model, because different decisions may be optimal depending on different ways they are valued (e.g., optimistically vs. pessimistically), which is encoded in the *Q*^−^ to *Q*^+^ interval. By contrast, exploration is used to account for sampling error in how decisions are valued, so that decisions can be considered that might be incorrectly valued.

In summary, risk-sensitivity parameters *k*^±^ can capture a range of behavior from being too risky to not risky enough and from too decisive to too deliberative. We demonstrate these decision-making behaviors in the next two examples.

### Capturing a penchant for gambling

To demonstrate how risk-sensitivity parameters drive decision-making behavior in our model, let us consider the IGT, which we introduced earlier. Selecting bad decks was put forth as a marker of impaired decision-making, or more specifically, an insensitivity to future consequences. This interpretation, however, presumes that the participant’s objective is indeed to make decisions that maximize expected rewards as opposed to making decisions that seeks large gains or avoids large losses. Risk-seeking behavior (i.e. a penchant for gambling), in particular, may encourage individuals to pursue bad decks, since they yield the largest one-time gains.

To that point, balanced (*k*^+^, *k*^−^) = (0.9, 0.9) risk-sensitivity parameters, reflecting risk-neutral behavior, results in a preference for Deck A, i.e. one of the good decks that leads to average net gains (Fig 6A). By contrast, imbalanced (*k*^+^,*k*^−^) = (0.9,0.1) risk-sensitivity parameters, reflecting risk-seeking behavior, results in a preference for Deck B, i.e. one of the bad decks that leads to average net losses. In each case, pessimistic state-action values *Q*^−^ are larger for good decks (C and D), correctly signifying that these decks are the more risk-averse choices (Fig 6B). Meanwhile, optimistic state-action values *Q*^+^ are larger for bad decks (A and B), correctly signifying that these decks are the more risk-seeking choices. Imbalanced risk-sensitivity parameters, however, dramatically underplays the risk of Deck B compared to balanced risk-sensitive parameters. Consequently, the chance of large gains encoded in *Q*^+^ is suitably enticing to encourage a Deck B preference. That is, Deck B preference, which is actually a well-known phenomenon of healthy participants [32], can be interpreted as a penchant for gambling rather than an insensitivity to future consequences.

**Fig 6.**
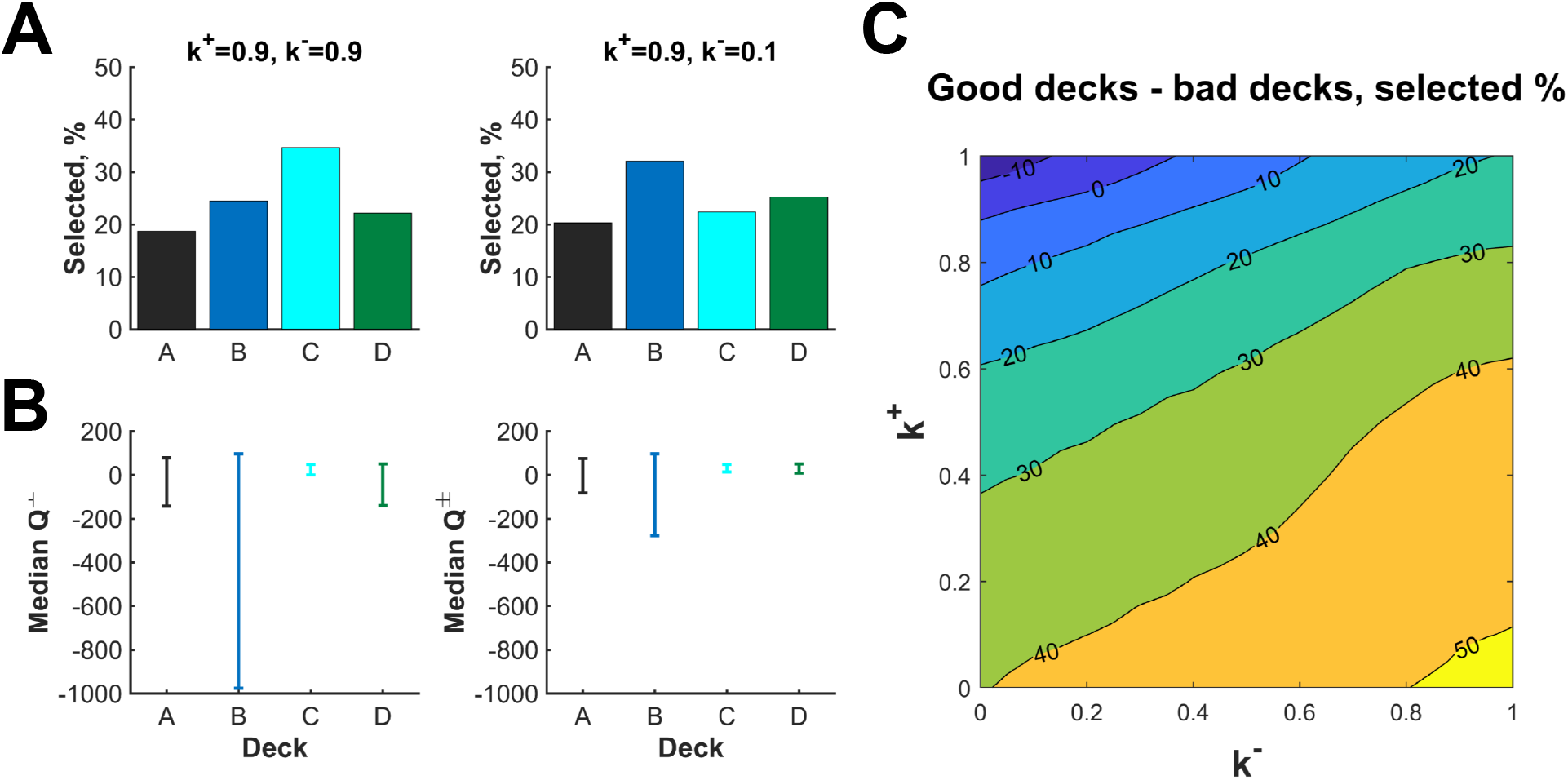
Risk-sensitivity of the Competing-Critics model during the Iowa Gambling task aggregated over 100 trials and 20,000 simulations. **A)** The “risky” Deck B becomes the most popular choice rather than Deck C, when parameter *k*^−^ is decreased from 0.9 to 0.1. **B)** Deck B becomes more favorable because of a dramatic increase to the pessimistic value function *Q*^−^. **C)** Bad decks A and B are chosen at higher rates moving along the risk-sensitivity axis (i.e. the *k*^+^ = 1 − *k*^−^ line).

As was done in [37], we can also partition the parameter space {(*k*^+^, *k*^−^)| 0 ≤ *k*^+^, *k*^−^ ≤ 1} by preference for good and bad decks (Fig 6C). This figure tells us that in the “blue” region of the parameter space, bad decks *A* + *B* are selected at greater frequency than good decks *C* + *D*. In the context of risk-seeking vs risk-averse terminology, our choice of *k*^+^ >> *k*^−^ means that our learner, despite the fact that B incurs incomparably large loss, keeps sticking to it because *Q*^+^ is driving the choice. In another words, our agent is unable to learn the good decks in the IGT, thus mimicking the behaviors of the participants with prefrontal cortex damage as demonstrated in [38].

### The ambiguity of deliberation

To demonstrate how parameters *k*^±^ drives uncertainty-sensitivity in the Competing-Critics model, let us consider the 2-stage Markov task, mentioned earlier. Recall this task uses the gap in stay probabilities to determine the degree to which individuals are learning about the common (*p* = 0.7) versus rare (*p* = 0.3) transition associated with each action in stage 1. As with the IGT, this interpretation supposes that individuals seeks to maximize excepted (discounted) rewards, rather than balance their internal optimist and pessimist. To account for the multi-stage aspect of this task, we use a discount factor *γ* to 0.9 which allows information to pass between task stages.

The bar graphs in Figure 7 represents the probability of sticking to the current choice categorized by whether it resulted in reward or not and whether the transition was common or rare. The contour plots on the right side of the figure tells us the difference between the probabilities of staying when the transition was common or rare, given rewarded or unrewarded.

**Fig 7.**
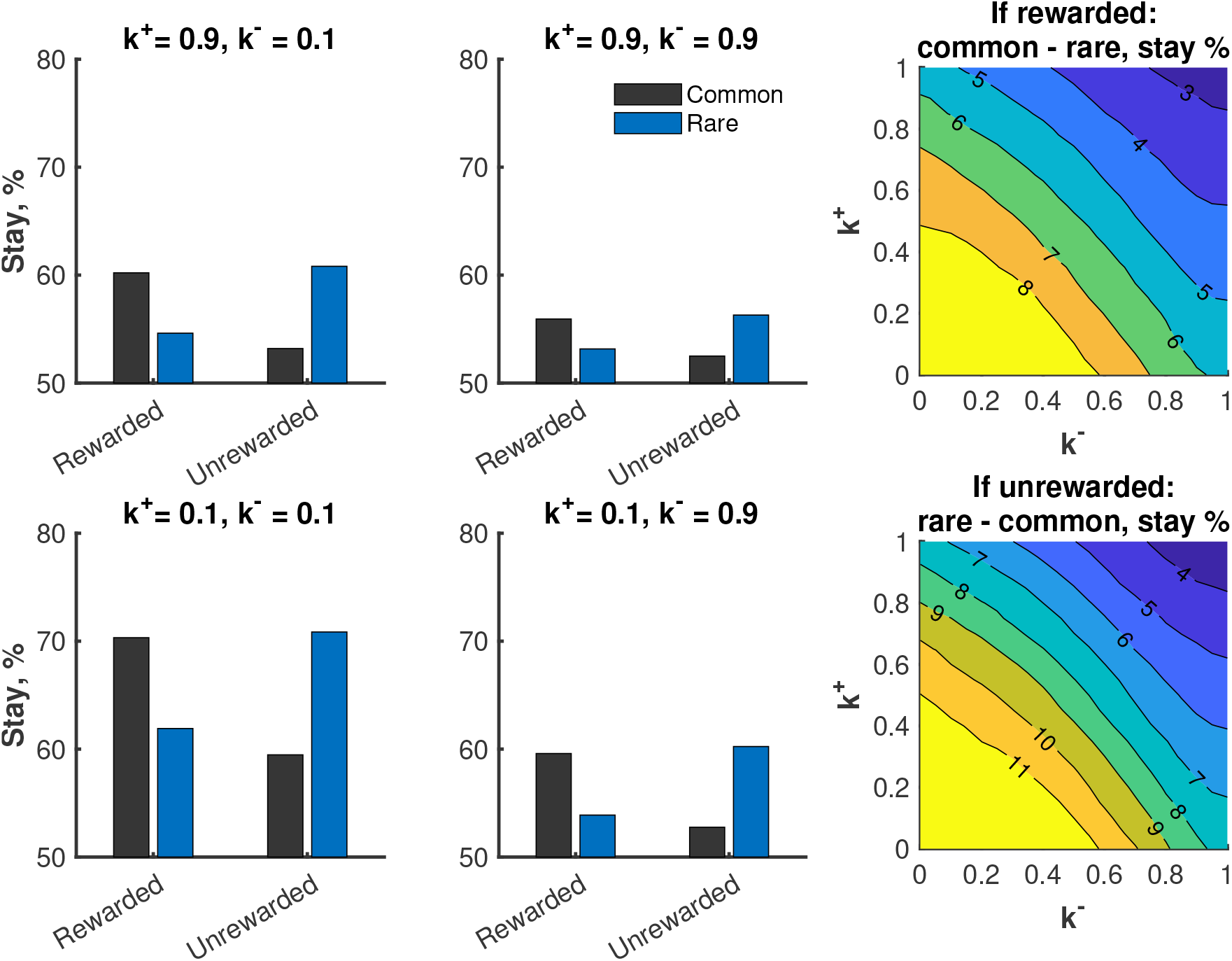
Stay probabilities after a first stage choice over a horizon of 80 decisions (40 first-stage decisions) and 20,000 simulations. The gap between stay probabilities for common vs. rarer transitions increases along the uncertainty-sensitivity axis (i.e. *k*^+^ = *k*^−^ axis) as the learner increases their deliberation about multiple choices.

For instance, whether the current decision was rewarded or not, the agent tends to become very deliberative if we let *k*^+^, *k*^−^ → 1. This case captures the behavior of someone who is both risk-seeking and risk-averse at the same time. Balancing these polar opposite behaviors is perhaps difficult, thus resulting in stagnated actions. One can see that in this case all 4 probabilities are roughly the same, meaning the individual does not discriminate between characteristics of the task.

On the other hand, our model stipulates that the choice of (*k*^+^, *k*^−^) = (0.1,0.1) should give an agent who is rather risk-neutral and behaves rather optimally with respect to expected rewards, because *k*^+^ = *k*^−^ = 0 will give the regular SARSA learner. One can see that the agent is more likely to stay when the current choice was rewarded. Furthermore, the agent is most likely to stay when the transition was common which led to a reward and least likely to stay when the transition was common which did not lead to a reward.

Regardless of the choice of (*k*^+^, *k*^−^), our model does not change the degree to which individuals are learning about common versus rare transitions. Yet, if we would only look at the gap in stay probabilities, we would conclude the smaller values of (*k*^+^, *k*^−^) are associated with better ability to learn these transitions. Further, a person’s tendency to explore decisions will also have a lower gap in stay probabilities. In other words, it is difficult to disambiguate a change in how deliberative a person is with their decisions from their ability to learn transitions or their tendency to explore.

### Predicting reaction time

One prediction of the Competing-Critics model is that the translation of *Q*^+^ and *Q*^−^ into decisions plays out in time. The value *Q*^+^ determines when (in time) a decision is considered, so that decisions with larger *Q*^+^ are considered earlier, and the value *Q*^−^ determines when a decision is no longer considered, so that decisions with smaller *Q*^−^ are considered for longer. Mathematically, we capture this idea by letting the decisions a compete for the largest random uniform draw *U_t_*(*a*) from (*Q*^−^(*S_t_, a*), *Q*^+^ (*S_t_, a*)). We propose that there is some non-increasing function that maps the max draw to time and that may depend on the person and the task. Larger max_*a*_ *U_t_*(*a*) leads to faster decisions; smaller to slower decisions.

In light of this prediction, we simulated max_*a*_ *U_t_*(*a*) for both the IGT and two-stage Markov task (Figure 8). In both tasks, larger max_a_ *U_t_*(*a*) are found for large *k*^+^ and small *k*^−^, i.e. when the overall behavior is more risk-seeking. Hence, our model predicts that risk-seeking decisions are quick and risk-averse decisions are slow.

**Fig 8.**
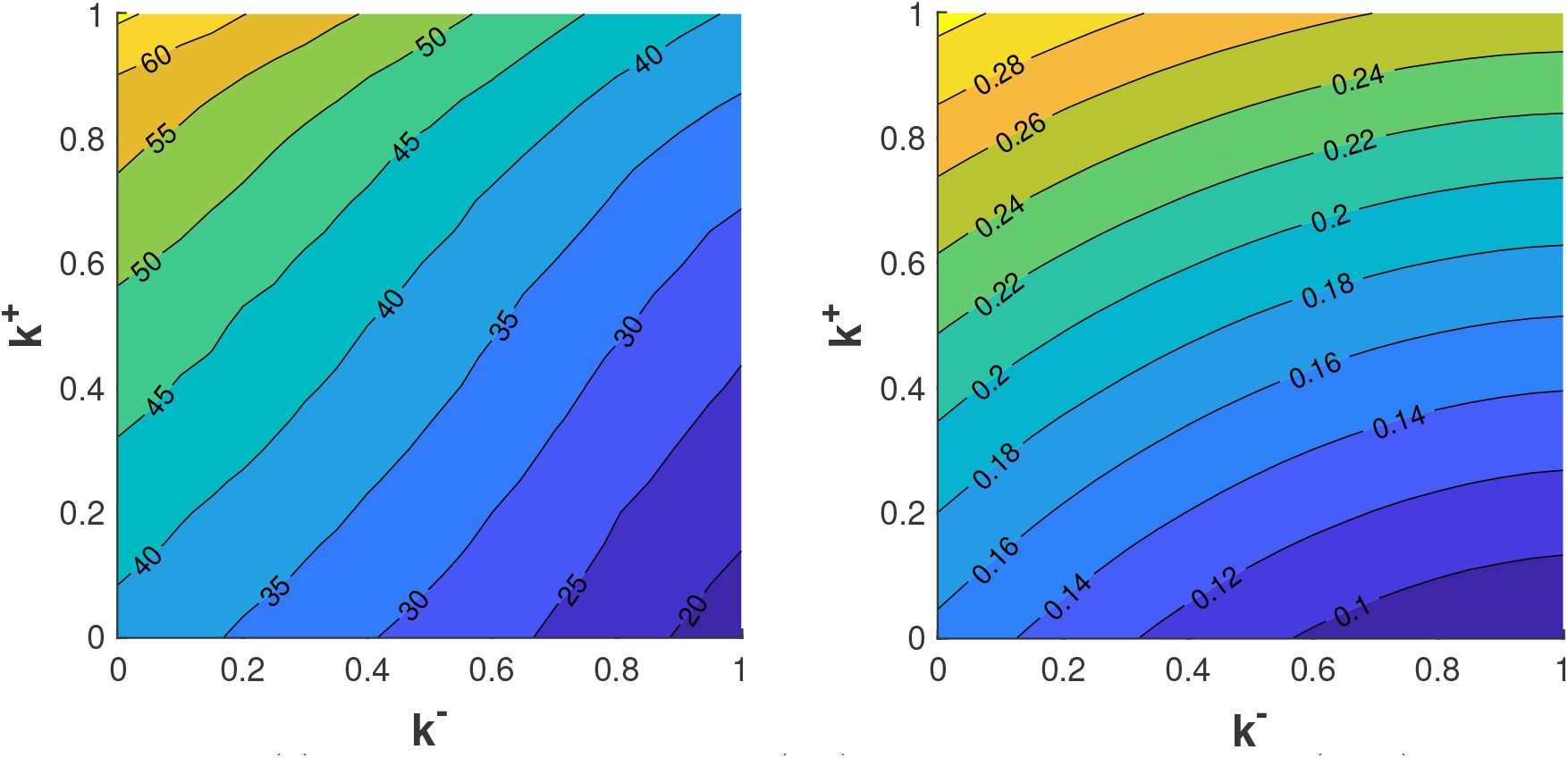
Mean max_*a*_ *U_t_*(*a*) in the Iowa Gambling Task (left) and two-stage Markov task (right). Larger values of max_*a*_ *U_t_*(*a*) are hypothesized to correspond to faster reaction times.

### Neural encoding of updates

As we mentioned, the rough intuition behind the reinforcement learning update we chose for the state-value functions *Q*^+^ and *Q*^−^ is that they capture the behaviors of risk-seeking and risk-averse learners, respectively. Going even further, we investigate the possibility that dopamine transients encode the update Δ*Q*^+^ associated with the risk-seeking system and serotonin transients encode the negative of the update Δ*Q*^−^ associated with the risk-averse system. In view of this claim, we present one last study, which measured dopamine and serotonin during a decision-making task [7].

In this study, participants were asked to make investing decisions on a virtual stock market. In total, participants made 20 investment decisions for 6 markets for a total of 120 decisions. Each participant was allocated $100 at the start of each market and could allocate bets between 0% to 100% in increments of 10%. The participant would gain or lose money depending on their bet. Given a bet *A_t_* on trial *t* and market value *p*_*t*+1_ after betting, percent monetary gain (or loss) on trial *t* was

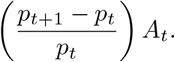

To model this experiment, we use the simplifying assumption that bets are low or high: *A_t_* = {25%, 75%}, and suppose rewards are

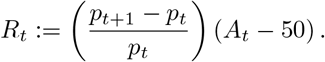

Actions are centered to 50% to account for the hypothesized role of counterfactual in this experiment [8]. Hence, *R_t_* is the percent monetary gain relative to the *counterfactual* gain were a neutral 50% bet made. Following [7], trials are split according to a reward prediction error (RPE): the percent monetary gain centered to its past mean and inversely scaled by its past standard deviation.

Let us consider the scenario where a decision made on trial *t* resulted in a negative RPE, which means that a lower monetary gain relative to past expected gains (Left panel in Figure 9). Without accounting for counterfactuals, a risk-neutral system would experience a negative update independent of bet level. Risk-seeking update Δ*Q*^+^, however, depends on bet level during a negative RPE: large for a low bet (25%) compared to a high bet (75%). The reverse is true for the negative of the risk-averse update Δ*Q*^−^: it is large for a high bet (75%) compared to a low bet (25%).

**Fig 9.**
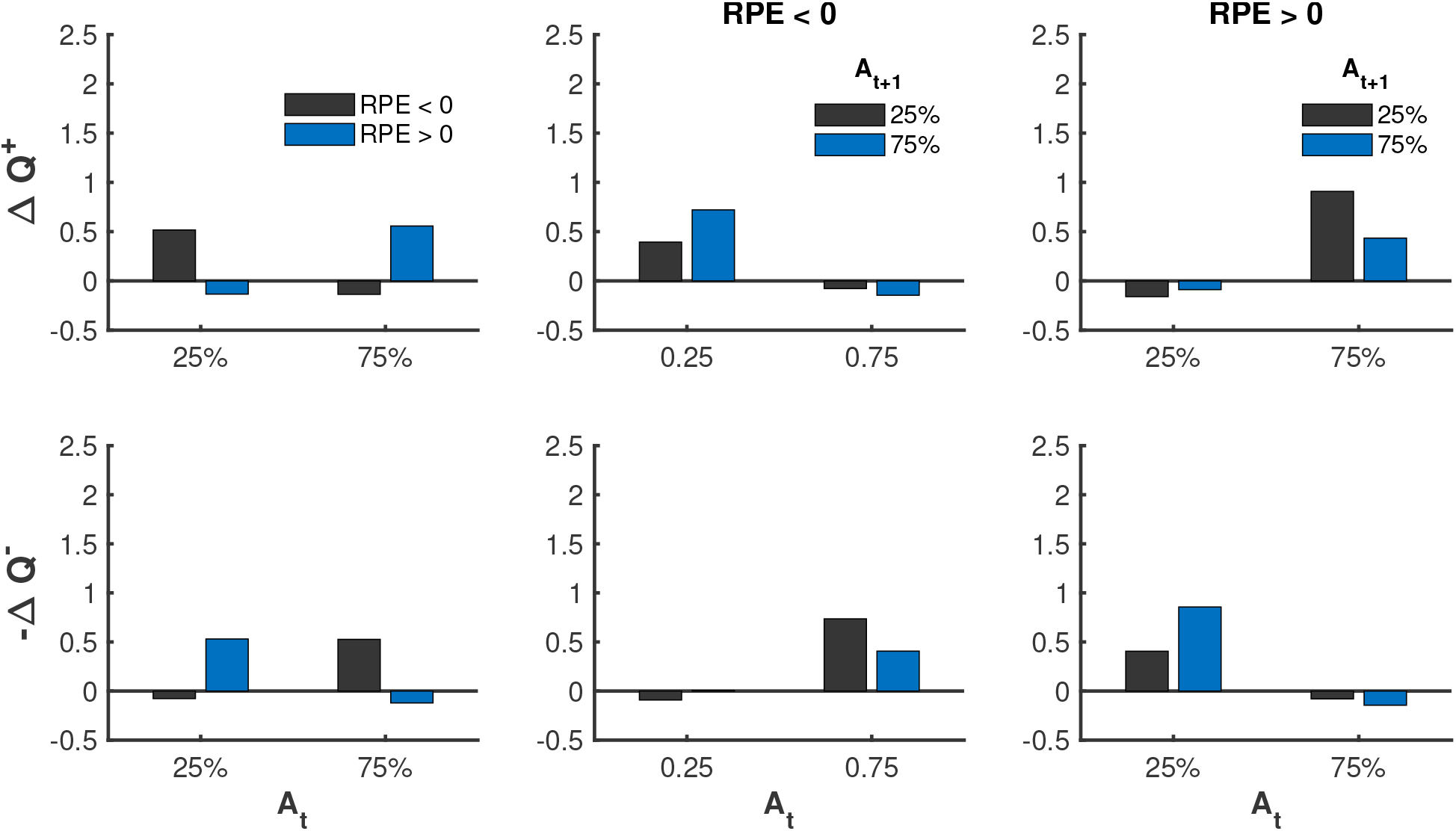
Mean updates as a function of bet levels and reward prediction error (RPE) over 20,000 simulations. Mirroring dopamine transients in [8], large mean Δ*Q*^+^ reinforces either a large bet for positive RPE or a small bet when negative RPE. Mirroring serotonin transients in [7], large mean −*Q*^−^ reinforces either a large bet for negative RPE or a small bet for positive RPE. In addition, mean updates can predict the upcoming bet and are asymmetrical, respecting potential asymmetry in the degree to which dopamine and serotonin transients can increase vs. decrease.

These characteristics of Δ*Q*^+^ and −*Q*^−^ during negative RPE mirror, respectively, dopamine and serotonin transients in [7]. The authors hypothesized that the large dopamine transient for a low bet encourages the *rewarding* decision of betting low, whereas the large serotonin transient for a high bet protects the individual from the *risky* decision of betting high. Betting low is only rewarding when compared to the counterfactual loss of betting a higher amount and losing. This hypothesis is consistent with the role in the Competing-Criticsmodel of a positive Δ*Q*^+^ to encourage a rewarding decision and a negative Δ*Q*^−^ to protect oneself from risky decisions.

When RPE is positive, which is when the agent has a lower monetary gain relative to past expected gains, the direction of the updates flip. The update Δ*Q*^+^ is now large for a high bet (75%) compared to a low bet (25%), and the negative of Δ*Q*^−^ is large for a low bet (25%) compared to a high bet (75%). Again, these characteristics mirror dopamine and serotonin transients in [7, 8]. In this case, it was hypothesized that the relatively large dopamine transient for a high bet encourages the rewarding decision of betting high, whereas the relatively large serotonin transient for a low bet protects the individual from the risky decision of betting low. As before, betting low is only considered risky when compared to the counterfactual loss of what they could have gained if they bet higher.

As an aside, we point out that average updates Δ*Q*^+^ and −*Q*^−^ are generally more positive than they are negative. This asymmetry respects the fact that dopamine and serotonin transients have a biophysical constraint whereby positive transients are easily induced but negative transients are not.

Following [7], we consider how updates Δ*Q*^+^ and −*Q*^−^ influence how a person subsequently bets (Middle and right panels in Figure 9). Trials are split further based on the subsequent decision made on the next trial. The negative of update Δ*Q*^−^ is largest when switching from a high to low bet during negative RPE and from a low to high bet during positive RPE. These trends mirror serotonin transients in [7], where a relatively large serotonin transient preceded a lowering of a bet when RPE was negative and preceded a raising or holding of a bet when RPE was positive. These findings provided further support that serotonin transients protect an individual from actual and counterfactual losses.

Meanwhile, the update Δ*Q*^+^ is largest when switching from a low to high bet during negative RPE and from a high to low bet during positive RPE. Since dopamine transients were not investigated as a function of subsequent bets in [7], we have the following model prediction without experimental validation: a relatively large dopamine transient preceding a raising of a bet when RPE was negative and preceding a lowering a bet when RPE was positive.

## Discussion

We presented a computational model of human decision-making called the Competing-Critics model. The model conceptualizes decision-making with two competing critics, an optimist and a pessimist, which are modulated by parameters *k*^+^ and *k*^−^, respectively. We posit that information is integrated from each system over time while decisions compete. The optimist activates decisions (“go”); the pessimist inhibits decisions (“no-go”). We show how our model can illuminate behavior observed in experiments using the Iowa gambling, two-stage Markov, or the stock market tasks.

A key prediction of the Competing-Critics model is that the updates in the optimistic and pessimistic learning systems are directly encoded in dopamine and serotonin transients. This finding arose from efforts to reproduce observations during the stock market task in Moran *et al* [7] and Kishida *et al* [8]. While computational models such as TD learning have provided a useful framework to interpret experiments involving dopamine [39], serotonin has been more difficult to pin down [19]. If serotonin can be understood as updates to a pessimistic learning system, then we would expect serotonin, like dopamine, to influence decision-making in important ways. It would oppose dopamine, protect a person from risky behavior, inhibit certain decisions, and change the value (and timing) of decisions. These functions agree with several leading theories [6, 7, 18–22]; yet, the mathematical form we propose for serotonin is new.

We are not the first to try to interpret observations of serotonin and dopamine through the lens of a computational model [6, 18, 30, 40]. Daw *et al*, for instance, describe how prediction error in a TD learning system could be transformed into tonic and phasic parts of dopamine and serotonin signals [6]. Alternatively, Montague *et al* argue that two prediction errors, derived from reward-predicting and aversive-predicting TD learning systems, could be transformed into serotonin and dopamine transients [18]. While these models map prediction errors to dopamine and serotonin, the more useful task might be mapping dopamine and serotonin to learning. In other words, trying to understand what certain dopamine and serotonin transients could mean to how a person learns and makes decisions. Our model provides a surprisingly simple answer: dopamine and serotonin transients are exactly the updates to two learning systems.

Critically, these learning systems can capture ranges of decision-making behavior. These learning systems (and hence, dopamine and serotonin) may oppose each other, but they are not perfect antipodes. Hence, the systems are not redundant and obey a principle about efficient coding of information [18]. For instance, we show that the two learning systems in the Competing-Critics model can implicitly reflect at least two properties of rewards: reward expectations and reward uncertainty. Several other mathematical models of learning and decision-making suggest individuals track reward uncertainty, but do so explicitly [10–14].

In addition, the Competing-Critics model reveals how risk-sensitivity and uncertainty-sensitivity represent two orthogonal dimensions of decision-making and how extreme values in either direction could pose unique impairments in decision-making. Sensitivity to risk and uncertainty are well documented in the psychological, economics, and reinforcement learning literature. For instance, risk-seeking (risk-aversion) can be beneficial when large rewards (small losses) are required to escape (avoid) bad scenarios. Platt provides several examples of animals behaving in a risk-sensitive way, e.g., birds switching from risk-aversion to risk-seeking as a function of the temperature [41]. Miscalibrated risk-sensitivity is thought to cause significant problems for people and underlie a number of psychiatric conditions such as addiction or depression [28, 29]. Mathematically, risk-sensitivity is captured either explicitly through functions that reflect risk-sensitive objectives [42, 43] or implicitly through differential weighting of positive and negative prediction errors [15–17], such as we do here. We recommend the paper by Mihatsch *et al* [27] for a nice theoretical treatment of risk-sensitivity.

Meanwhile, uncertainty-sensitivity represents the degree to which uncertainty in the reward distribution, and in their knowledge of this distribution, influences their decisions. Like risk-sensitivity, miscalibrated uncertainty-sensitivity is thought to underlie psychiatric conditions such as anxiety [44–47]. Huang *et al*, for example, describe this miscalibration in anxiety as a “failure to differentiate signal from noise” leading to a “sub-optimal” decision strategy [45]. Conceptually, our model provides a different interpretation. Rather than being a failure or sub-optimal behavior, extreme uncertainty-sensitivity embodies a strategy that attempts to satisfy competing objectives, some of which are risk-averse and others which are risk-seeking. In experiments, this conflicted strategy will look similar to an exploration-exploitation trade-off, making it difficult to distinguish between the two.

Interestingly, any attempt to modify solely the optimistic and pessimistic learning system (or dopamine and serotonin transients) will affect both risk sensitivity and uncertainty-sensitivity. The reason is that risk-sensitivity and uncertainty-sensitivity axes are rotated 45 degrees from the axes of the parameters *k*^+^ and *k*^−^ modulating the two learning systems. For instance, increasing *k*^−^ in an attempt to reduce risk-seeking would have the unintended consequence of increasing the sensitivity to uncertainty. Under our interpretation, this would correspond to interventions on serotonin transients to reduce risk-seeking having the potential side-effect of a loss of decisiveness. Similarly, reducing *k*^+^, or intervening on dopamine transients, to reduce risk-seeking would decrease sensitivity to uncertainty. A similar tradeoff occurs when trying to decrease risk-aversion or sensitivity to uncertainty through manipulations of just *k*^+^ or just *k*^−^. Notably, many current pharmacological interventions (e.g., Lithium) act on both dopamine and serotonin neurons.

Another key prediction of our model is that values placed on decisions by the two learning system (i.e. *Q*^±^) determine the time to make a decision. Thus, the distribution of reaction time may provide additional data beyond choice selection for which to inform or falsify our model. This connection to reaction time might also help to make sense of the impact of serotonin and dopamine on how quickly decisions are made (e.g., impulsively) [19, 26, 48]. Models for reaction time are often built with stochastic differential equations such as drift-diffusion models to reflect a process of evidence accumulation (c.f., [49, 50] for an overview). For example, drift-diffusion models of reaction time can be integrated with a TD learning model by relating drift velocities to different in values between two choices [51]. Reaction time in our model differs from this approach in that it can arise from any number of possible decisions, as opposed to just two, and is sensitive to risk and uncertainty, rather than a single value, for each decision. This additional flexibility may be useful for explaining experimental observations of reaction time.

There are several limitations of this work to consider. We hope it is clear that the modeling of learning in the updates of *Q*^+^ and *Q*^−^ is largely modular from the modeling that maps these values to actions and reaction times. There are numerous ways that pairs of *Q*^+^ and *Q*^−^ values can be mapped to a choice of actions and a time delay in making that choice. In addition, our model was built upon a SARSA algorithm, but Q-learning may prove to be equally suitable. It should also be clear that our model is over-simplified. One notable absence, for example, is that our model did not track average outcomes or map these outcomes or other parts of our model to tonic dopamine and serotonin, unlike the model of Daw *et al* [6]. Relatedly, we directly incorporated counterfactuals into our rewards to reproduce findings from the stock market task [7, 8], but perhaps a separate process, such as tonic serotonin or dopamine, should be included to track counterfactuals. Another limitation of our model is that it relies on only two prediction errors. However, a recent study suggests dopamine is capable of capturing a distribution of prediction errors [9], which has the advantage of being able to learn about the distribution of rewards.

Lastly, one of the key properties of our model, the ordering 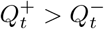 assumes that the parameter *α* is the same for *Q*^+^ and *Q*^−^. If parameter *α* were not equal, then the relationship between *Q*^+^ and *Q*^−^ could reverse. The possible effects of *Q*^+^ < *Q*^−^ largely fall outside the specifics of the Competing-Critics model, but it is conceivable such a situation could result in no-go signals arriving before go signals, leading to a decision process unwilling to even consider an option. A situation when no options were even worth consideration may be similar to anhedonia.

In conclusion, this work establishes a new model of human decision-making to help illuminate, clarify, and extend current experiments and theories. Such a model could be utilized to quantify normative and pathological ranges of risk-sensitivity and uncertainty-sensitivity. Overall, this work moves us closer to a precise and mechanistic understanding of how humans make decisions.

## Author Contributions

All authors contributed to the initial conceptualization of the model. EE wrote the simulation code, conducted the simulation experiments, and derived mathematical properties of the model. EE and ALC produced the figures. EE, ALC, and JN drafted different sections of the paper. All authors reviewed and edited the paper.

## Notes

### Competing Interest Statement

The authors have declared no competing interest.

### Summary of Updates

We have updated the model name and paper title, and added a few sentences. A new section has been added to the supplement.

## References

1. Schultz W, Dayan P, Montague PR. A neural substrate of prediction and reward. Science. 1997;275(5306):1593–1599.

2. Schultz W, Apicella P, Ljungberg T. Responses of monkey dopamine neurons to reward and conditioned stimuli during successive steps of learning a delayed response task. Journal of neuroscience. 1993;13(3):900–913.

3. Zaghloul KA, Blanco JA, Weidemann CT, McGill K, Jaggi JL, Baltuch GH, et al. Human substantia nigra neurons encode unexpected financial rewards. Science. 2009;323(5920):1496–1499.

4. Pan WX, Schmidt R, Wickens JR, Hyland BI. Dopamine cells respond to predicted events during classical conditioning: evidence for eligibility traces in the reward-learning network. Journal of Neuroscience. 2005;25(26):6235–6242.

5. Cohen JY, Haesler S, Vong L, Lowell BB, Uchida N. Neuron-type-specific signals for reward and punishment in the ventral tegmental area. nature. 2012;482(7383):85.

6. Daw ND, Kakade S, Dayan P. Opponent interactions between serotonin and dopamine. Neural Networks. 2002;15(4-6):603–616.

7. Moran RJ, Kishida KT, Lohrenz T, Saez I, Laxton AW, Witcher MR, et al. The protective action encoding of serotonin transients in the human brain. Neuropsychopharmacology. 2018;43(6):1425.

8. Kishida KT, Saez I, Lohrenz T, Witcher MR, Laxton AW, Tatter SB, et al. Subsecond dopamine fluctuations in human striatum encode superposed error signals about actual and counterfactual reward. Proceedings of the National Academy of Sciences. 2016;113(1):200–205.

9. Dabney W, Kurth-Nelson Z, Uchida N, Starkweather CK, Hassabis D, Munos R, et al. A distributional code for value in dopamine-based reinforcement learning. Nature. 2020;577(7792):671–675.

10. Li J, Schiller D, Schoenbaum G, Phelps EA, Daw ND. Differential roles of human striatum and amygdala in associative learning. Nature neuroscience. 2011;14(10):1250.

11. Jepma M, Schaaf JV, Visser I, Huizenga HM. Uncertainty-driven regulation of learning and exploration in adolescents: A computational account. PLoS computational biology. 2020;16(9):e1008276.

12. Redish AD, Jensen S, Johnson A, Kurth-Nelson Z. Reconciling reinforcement learning models with behavioral extinction and renewal: implications for addiction, relapse, and problem gambling. Psychological review. 2007;114(3):784.

13. Gershman SJ, Monfils MH, Norman KA, Niv Y. The computational nature of memory modification. Elife. 2017;6:e23763.

14. Angela JY, Dayan P. Uncertainty, neuromodulation, and attention. Neuron. 2005;46(4):681–692.

15. Ross MC, Lenow JK, Kilts CD, Cisler JM. Altered neural encoding of prediction errors in assault-related posttraumatic stress disorder. Journal of psychiatric research. 2018;103:83–90.

16. Hauser TU, Iannaccone R, Walitza S, Brandeis D, Brem S. Cognitive flexibility in adolescence: neural and behavioral mechanisms of reward prediction error processing in adaptive decision making during development. Neuroimage. 2015;104:347–354.

17. Niv Y, Edlund JA, Dayan P, O’Doherty JP. Neural prediction errors reveal a risk-sensitive reinforcement-learning process in the human brain. Journal of Neuroscience. 2012;32(2):551–562.

18. Montague PR, Kishida KT, Moran RJ, Lohrenz TM. An efficiency framework for valence processing systems inspired by soft cross-wiring. Current opinion in behavioral sciences. 2016;11:121–129.

19. Cools R, Nakamura K, Daw ND. Serotonin and dopamine: unifying affective, activational, and decision functions. Neuropsychopharmacology. 2011;36(1):98–113.

20. Deakin JW, Graeff FG. 5-HT and mechanisms of defence. Journal of psychopharmacology. 1991;5(4):305–315.

21. Deakin J. Roles of serotonergic systems in escape, avoidance and other behaviours. Theory in psychopharmacology. 1983;2:149–193.

22. Rogers RD. The roles of dopamine and serotonin in decision making: evidence from pharmacological experiments in humans. Neuropsychopharmacology. 2011;36(1):114–132.

23. Sutton RS, Barto AG. Reinforcement learning: An introduction. MIT press; 2018.

24. Rescorla RA, Wagner AR, et al. A theory of Pavlovian conditioning: Variations in the effectiveness of reinforcement and nonreinforcement. Classical conditioning II: Current research and theory. 1972;2:64–99.

25. Bayer HM, Glimcher PW. Midbrain dopamine neurons encode a quantitative reward prediction error signal. Neuron. 2005;47(1):129–141.

26. Niv Y, Duff MO, Dayan P. Dopamine, uncertainty and TD learning. Behavioral and brain Functions. 2005;1(1):6.

27. Mihatsch O, Neuneier R. Risk-sensitive reinforcement learning. Machine learning. 2002;49(2-3):267–290.

28. Korn C, Sharot T, Walter H, Heekeren H, Dolan RJ. Depression is related to an absence of optimistically biased belief updating about future life events. Psychological medicine. 2014;44(3):579–592.

29. Rouhani N, Niv Y. Depressive symptoms bias the prediction-error enhancement of memory towards negative events in reinforcement learning. Psychopharmacology. 2019;236(8):2425–2435.

30. Dayan P, Huys QJ. Serotonin, inhibition, and negative mood. PLoS Comput Biol. 2008;4(2):e4.

31. Dayan P, Huys QJ. Serotonin in affective control. Annual review of neuroscience. 2009;32.

32. Chiu YC, Huang JT, Duann JR, Lin CH. Twenty years after the iowa gambling task: rationality, emotion, and decision-making. Frontiers in psychology. 2018;8:2353.

33. Bechara A, Damasio AR, Damasio H, Anderson SW, et al. Insensitivity to future consequences following damage to human prefrontal cortex. Cognition. 1994;50:1–3.

34. Daw ND, Gershman SJ, Seymour B, Dayan P, Dolan RJ. Model-based influences on humans’ choices and striatal prediction errors. Neuron. 2011;69(6):1204–1215.

35. Collins AG, Frank MJ. Opponent actor learning (OpAL): Modeling interactive effects of striatal dopamine on reinforcement learning and choice incentive. Psychological review. 2014;121(3):337.

36. Mikhael JG, Bogacz R. Learning reward uncertainty in the basal ganglia. PLoS computational biology. 2016;12(9):e1005062.

37. Steingroever H, Wetzels R, Wagenmakers EJ. A Comparison of Reinforcement Learning Models for the Iowa Gambling Task Using Parameter Space Partitioning. Journal of Problem Solving. 2013;5(2).

38. Lin CH, Chiu YC, Lee PL, Hsieh JC. Is deck B a disadvantageous deck in the Iowa Gambling Task? Behavioral and Brain Functions. 2007 Mar;3(1):16. Available from: https://doi.org/10.1186/1744-9081-3-16.

39. Glimcher PW. Understanding dopamine and reinforcement learning: the dopamine reward prediction error hypothesis. Proceedings of the National Academy of Sciences. 2011;108(Supplement 3):15647–15654.

40. Priyadharsini BP, Ravindran B, Chakravarthy VS. Understanding the role of serotonin in basal ganglia through a unified model. In: International Conference on Artificial Neural Networks. Springer; 2012. p. 467–473.

41. Platt ML, Huettel SA. Risky business: the neuroeconomics of decision making under uncertainty. Nature neuroscience. 2008;11(4):398.

42. Kahneman D, Tversky A. Choices, values, and frames. In: Handbook of the fundamentals of financial decision making: Part I. World Scientific; 2013. p. 269–278.

43. Glimcher PW, Rustichini A. Neuroeconomics: the consilience of brain and decision. Science. 2004;306(5695):447–452.

44. Hirsh JB, Mar RA, Peterson JB. Psychological entropy: A framework for understanding uncertainty-related anxiety. Psychological review. 2012;119(2):304.

45. Huang H, Thompson W, Paulus MP. Computational dysfunctions in anxiety: Failure to differentiate signal from noise. Biological psychiatry. 2017;82(6):440–446.

46. Grupe DW, Nitschke JB. Uncertainty and anticipation in anxiety: an integrated neurobiological and psychological perspective. Nature Reviews Neuroscience. 2013;14(7):488–501.

47. Luhmann CC, Ishida K, Hajcak G. Intolerance of uncertainty and decisions about delayed, probabilistic rewards. Behavior Therapy. 2011;42(3):378–386.

48. Worbe Y, Savulich G, Voon V, Fernandez-Egea E, Robbins TW. Serotonin depletion induces ‘waiting impulsivity’ on the human four-choice serial reaction time task: cross-species translational significance. Neuropsychopharmacology. 2014;39(6):1519–1526.

49. Kilpatrick ZP, Holmes WR, Eissa TL, Josie K. Optimal models of decision-making in dynamic environments. Current Opinion in Neurobiology. 2019;58:54–60.

50. Veliz-Cuba A, Kilpatrick ZP, Josic K. Stochastic models of evidence accumulation in changing environments. SIAM Review. 2016;58(2):264–289.

51. Pedersen ML, Frank MJ, Biele G. The drift diffusion model as the choice rule in reinforcement learning. Psychonomic bulletin & review. 2017;24(4):1234–1251.

