## Supplement for "A competition of critics in human decision-making"

### Stochastic Dominance

Let us index $Q^{\pm}$ by time $t$ so that the original update equation will look as follows:

$$Q^{\pm}_{t+1}(S_t, A_t) = Q^{\pm}_t(S_t, A_t) + \alpha \left[(1 \pm k^{\pm})\mathbb{1}_{\delta^{\pm}_t > 0} + (1 \mp k^{\pm})\mathbb{1}_{\delta^{\pm}_t < 0}\right] \delta^{\pm}_t$$

with $Q^{\pm}_0 = 0$ and

$$\delta^{\pm}_t = R_{t+1} + \gamma Q^{\pm}_t(S_{t+1}, A_{t+1}) - Q^{\pm}_t(S_t, A_t).$$

**Proposition 1.** *If $k^{\pm} \geq 0$ and $\alpha(1 - k^{\pm}) \leq 1$, then*

$$Q^{+}_t(s, a) \geq Q^{-}_t(s, a)$$

*for all pairs $(s, a) \in \mathcal{S} \times \mathcal{A}$ and all non-negative integers $t$.*

*Proof.* Assume $k^{\pm} \geq 0$ and $\alpha(1 - k^{\pm}) \leq 1$. Our proof is by induction. The base case $t = 0$ is trivial since $Q^{\pm}_0 = 0$. For clarity, let's compute $Q_1$. In this case, $\delta^{\pm}_1 = R_0$ and so

$$Q^{+}_1(S_0, A_0) = \alpha(1 + k^{+}\text{sign}(R_0))R_0$$

$$Q^{-}_1(S_0, A_0) = \alpha(1 - k^{-}\text{sign}(R_0))R_0.$$

Hence,

$$(Q_1^+ - Q_1^-)(S_0, A_0) = \alpha(k^+ + k^-)|R_0| \geq 0.$$

Now for the inductive hypothesis - assume that $Q_t^+(s,a) \geq Q_t^-(s,a)$ for all pairs $(s,a) \in \mathcal{S} \times \mathcal{A}$ and all non-negative integers up $t$. One can subtract $Q_{t+1}^+$ from $Q_{t+1}^-$ to obtain:

$$\begin{aligned}\left(Q^+ - Q^-\right)_{t+1}(S_t, A_t) = &\left(Q^+ - Q^-\right)_t(S_t, A_t) \\ &+ \alpha\left[(1+k^+)\mathbb{1}_{\delta_t^+ > 0} + (1-k^+)\mathbb{1}_{\delta_t^+ < 0}\right]\delta_t^+ \\ &- \alpha\left[(1-k^-)\mathbb{1}_{\delta_t^- > 0} + (1+k^-)\mathbb{1}_{\delta_t^- < 0}\right]\delta_t^-.\end{aligned}$$

Rewrite the above equation in the following form:

$$\begin{aligned}\left(Q^+ - Q^-\right)_{t+1}(S_t, A_t) = &(Q^+ - Q^-)_t(S_t, A_t) \\ &+ \alpha\mathbb{1}_{\{\delta_t^+ > 0 \geq \delta_t^-\}}\left[(1+k^+)\delta_t^+ - (1+k^-)\delta_t^-\right] \\ &+ \alpha\mathbb{1}_{\{\delta_t^+, \delta_t^- > 0\}}\left[(1+k^+)\delta_t^+ - (1-k^-)\delta_t^-\right] \\ &+ \alpha\mathbb{1}_{\{\delta_t^+, \delta_t^- \leq 0\}}\left[(1-k^+)\delta_t^+ - (1+k^-)\delta_t^-\right] \\ &+ \alpha\mathbb{1}_{\{\delta_t^+ \leq 0 < \delta_t^-\}}\left[(1-k^+)\delta_t^+ - (1-k^-)\delta_t^-\right]\end{aligned} \tag{1}$$

So it suffices to check that (1) is non-negative for the four cases for $\delta_t^-, \delta_t^+$.

**Case 1:** $\delta_t^+ > 0 \geq \delta_t^-$. In this case, (1) is trivially non-negative given the non-negativity of $k^\pm$ and the inductive assumption on $t$.

**Case 2:** $\delta_t^\pm > 0$. In this case, (1) is a linear function, $L(R_t)$, of $R_t$:

$$\begin{aligned}L(R_t) = &\left(Q^+ - Q^-\right)_t(S_t, A_t) \\ &+ \alpha(k^+ + k^-)R_t \\ &+ \alpha\gamma\left((1+k^+)Q_t^+(S_{t+1}, A_{t+1}) - (1-k^-)Q_t^-(S_{t+1}, A_{t+1})\right) \\ &- \alpha(1+k^+)Q_t^+(S_t, A_t) + \alpha(1-k^-)Q_t^-(S_t, A_t)\end{aligned}$$

Since $L$ has a non-negative slope $\alpha(k^+ + k^-)$, it stays non-negative if and only if it is non-negative on the lower bound of its domain. In particular, $\delta_t^\pm > 0$ gives lower bounds for $R_t$:

$$R_t > Q^\pm(S_t, A_t) - \gamma Q^\pm(S_{t+1}, A_{t+1}).$$

If $R_t = Q^+(S_t, A_t) - \gamma Q^+(S_{t+1}, A_{t+1})$, then $\delta_t^+ = 0$ and

$$\begin{aligned} L(R_t) &= (Q^+ - Q^-)_t(S_t, A_t) - \alpha(1 - k^-)\delta_t^- \\ &= (1 - \alpha(1 - k^-))(Q^+ - Q^-)_t(S_t, A_t) + \alpha\gamma(1 - k^-)(Q^+ - Q^-)_t(S_{t+1}, A_{t+1}) \geq 0 \end{aligned}$$

because of the conditions on $k^-$ and the inductive hypothesis. If $R_t = Q^-(S_t, A_t) - \gamma Q^-(S_{t+1}, A_{t+1})$, then $\delta_t^- = 0$ and

$$L(R_t) = (Q^+ - Q^-)_t(S_t, A_t) + \alpha(1 + k^+)\delta_t^+ \geq 0$$

because of $\delta_t^+ > 0$ and the inductive hypothesis.

**Case 3:** $\delta_t^+, \delta_t^- \leq 0$. Continuing along the similar line of reasoning, one can see that (1) is a linear function, $L(R_t)$, of $R_t$ with the non-positive slope $-\alpha(k^+ + k^-)$. Therefore, it suffices to check that $L$ is non-negative if and only if it is non-negative at its upper bound. Upper bounds for $R_t$ follow from $\delta_t^+, \delta_t^- \leq 0:$

$$R_t \leq Q_t^{\pm}(S_t, A_t) - \gamma Q_t^{\pm}(S_{t+1}, A_{t+1}).$$

If $R_t = Q_t^-(S_t, A_t) - \gamma Q_t^-(S_{t+1}, A_{t+1})$, then $\delta_t^- = 0$ and

$$\begin{aligned} L(R_t) &= (Q^+ - Q^-)_t(S_t, A_t) + \alpha(1 - k^+)\delta_t^+ \\ &= (1 - \alpha(1 - k^+))\left(Q^+ - Q^-\right)_t(S_t, A_t) + \alpha\gamma(1 - k^+)\left(Q^+ - Q^-\right)_t(S_{t+1}, A_{t+1}) \geq 0 \end{aligned}$$

because of conditions on $k^+$ and the inductive hypothesis. If $R_t = Q_t^+(S_t, A_t) - \gamma Q_t^+(S_{t+1}, A_{t+1})$, then $\delta_t^+ = 0$ and

$$L(R_t) = (Q^+ - Q^-)_t(S_t, A_t) - \alpha(1 + k^-)\delta_t^- \geq 0$$

immediately due to $\delta_t^- \leq 0$ and the inductive hypothesis.

**Case 4:** $\delta_t^+ \leq 0 < \delta_t^-$. This is the hardest case. Observe that (1) is again a linear function, $L(R_t)$, of $R_t$. Upper and lower bounds on $R_t$ can be derived from $\delta_t^+ \leq 0 < \delta_t^-$, or more explicitly:

$$Q_t^-(S_t, A_t) - \gamma Q^-(S_{t+1}, A_{t+1}) \leq R_t \leq Q_t^+(S_t, A_t) - \gamma Q_t^+(S_{t+1}, A_{t+1}).$$

Therefore, if $L(R_t)$ is non-negative when evaluated at these bounds, it is non-negative on its domain. At the lower bound, one has $\delta_t^- = 0$ and

$$\begin{aligned} L(R_t) &= (Q^+ - Q^-)_t(S_t, A_t) + \alpha(1-k^+)\delta_t^+ \\ &= (1-\alpha(1-k^+))(Q^+ - Q^-)_t(S_t, A_t) + \alpha\gamma(1-k^+)(Q^+ - Q^-)_t(S_{t+1}, A_{t+1}) \geq 0, \end{aligned}$$

which follows from conditions on $k^+$ and the inductive hypothesis. At the upper bound, one has $\delta_t^+ = 0$ and

$$\begin{aligned} L(R_t) &= (Q^+ - Q^-)(S_t, A_t) - \alpha(1-k^-)\delta_t^- \\ &= (1-\alpha(1-k^-))(Q^+ - Q^-)_t(S_t, A_t) + \alpha\gamma(1-k^-)(Q^+ - Q^-)_t(S_{t+1}, A_{t+1}) \geq 0, \end{aligned}$$

which follows from conditions on $k^\pm$ and the inductive hypothesis.

Thus, by induction, we obtain:

$$Q_t^+(s,a) \geq Q_t^-(s,a),$$

for all pairs $(s,a) \in \mathcal{S} \times \mathcal{A}$ and all non-negative integers $t$. □

We briefly remark about convergence of $Q^+$ and $Q^-$. While the authors expect that $Q^\pm$ converge in distribution under the same step-size condition in preceding proposition, a rigorous proof lies outside the scope of this section. Despite the fact that our model is based heavily on the classical reinforcement learning algorithm SARSA, the assumption of constant step-size make it difficult to apply the standard techniques of contraction-mapping used in standard literatures. Both in [1] and [2], one can see that it is crucial to have the condition:

1. $\sum_{t=0}^{\infty} \alpha_t = \infty$
2. $\sum_{t=0}^{\infty} \alpha_t^2 < \infty$

On the other hand, Theory of Iterated Random Functions deal with stochastic iterations with constant step-size. However, this usually comes at the expense of much stricter requirements on the rewards such as i.i.d $\{R_t\}$. Classic references on this subject are [3] and [4].

### Python Implementation

Here, we elaborate on the implementation of simulations done in the paper. Since we maintain two tables $Q^\pm$, two additional risk-parameters $k^\pm$, and a more elaborate action-selection method compared to the traditional SARSA algorithm, we give the skeleton of our python code below:

```python
# Defining the different parameters
epsilon = 0.3
S = 10000
T = 100
alpha = 0.5
gamma = 0.0

# In addition to the classical SARSA parameters
# we have k^+, k^-
k_plus = 0.9
k_minus = 0.9

#Initializing the Q^+, Q^- matrices
Q_plus = np.zeros((m,n))
Q_minus = np.zeros((m,n))

# helper function to choose action according to the Balanced-Sarsa
def choose_action(Q_plus, Q_minus, eps, curr_state):
    # CODE HERE
    ...
    return action

# helper function to update the Q-values
def update (curr_action, curr_state, Q, next_action, next_state, r,k,Q_type):
    # CODE HERE
    ...
# the function to encode the piecewise linear update
def piecewise(TD_error, k, Q_type):
    if Q_type == 'POS':
        if TD_error >= 0:
            return (1+k)*TD_error
        else:
            return (1-k)*TD_error
    elif TD_error >= 0:
        return (1-k)*TD_error
    else:
        return (1+k)*TD_error

# function to take a step in the environment
def step(curr_action, curr_state):
```

```
    # CODE HERE
    ...
    return reward, next_state

#---------Training here---------
for s in range(S)
    # Initialize the initial state and Q-tables here

    curr_action = choose_action(Q_plus, Q_minus, epsilon, curr_state)

    for t in range(T):
        reward, next_state = step(curr_state, curr_action)
        # select the "look-up" action as per SARSA
        next_action = choose_action(Q_plus, Q_minus, eps)

        # update the Q-values
        update(curr_action, curr_state,Q_plus, next_action, next_state, reward , k_plus, "POS")
        update(curr_action, curr_state, Q_minus, next_action, next_state, reward, k_minus, "NEG")

        curr_action = next_action
        curr_state = next_state
```

First of all, the functions **choose action**() , **update**(), and **piecewise**() are identical in all of the tasks, since they are only dependent on the algorithm.

On the other hand, the **step()** function is specific to each task since it needs to capture the transition probabilities and is thus implemented differently for each task. The full implementations can be found on: https://github.com/eza0107/Opposite-Systems-for-Decision-Making

### Iowa Gambling Task

In the original Iowa Gambling Task, the rewards followed a fixed, discrete distribution and was reset every 10 trials. For instance for Deck $A$, the first 10 trials included exactly 5 losses each worth \$250 while every trial also gave \$100 and the 5 losses were random within the 10 trial block. Our implementation modified this by choosing a binary reward from $\{\$100, -\$150\}$ with each option being equally likely for Deck $A$. This simplifies the reward structure a bit while preserving the loss-frequency and the expected gain over trials.

### Two-Stage MDP

For this task, complying with the original study in [5], the rewards received at the end of second-stage choice were altered by Gaussian noise with zero mean and 0.025 standard deviation at the end of each second-stage. On the other hand, the probability transitions between first-stage and second-stage were fixed at $\{\text{common, rare}\} = \{p = 0.7, p = 0.3\}$. The code snipper illustrates this below:

```
def reward_prob(T):
    q = np.zeros((2,2,T))
    q[0:2,0:2,0] = np.array([[0.75,0.75],[0.25,0.25]])

    for t in range(1,T):
        q[:,:,t] = q[:,:,t-1] + np.random.normal(0,0.025,[2,2])
        for i in range(2):
            for j in range(2):
                if q[i,j,t]>=1:
                    q[i,j,t] = 1 - (q[i,j,t] - 1)
                elif q[i,j,t]<=0:
                    q[i,j,t] = -q[i,j,t]
    return q
```

### Investment Task

Much like the IGT, we have one, stationary state and two actions here. The only point of importance when it comes to implementation is that we considered the counterfactual error and centered rewards, as mentioned previously:

```
def step(curr_action, curr_state, t, market_change):
    reward = (50*curr_action-25)*market_change[t]
    next_state = curr_state
    return reward, next_state
```

### Relation to Opposing Actor Learning (OpAL) model

The OpAL model in [6] is similar to our model in that it uses dual competing learning systems and relates these systems to “go” and ”no-go” signals. Despite these similarities, there are many other meaningful differences between the two models, which we expand on below. Let us first introduce the OpAL model. The OpAL model uses a single critic to place a value on each action $V_t(a)$. This value is updated for the selected action $A_t$ according to

$$V_t(A_t) = V_t(A_t) + \alpha\delta_t,$$

where

$$\delta_t = R_t - V_t(A_t).$$

We point out that these values do not depend on the state $S_t$ to keep with the original presentation of the OpAL model, but could be easily extended to account for state transitions. In the OpAL model, prediction error $\delta_t$ influences two populations of neurons, or “actors”, downstream. These populations are denoted by G to represent a go signal and N to represent a no-go signal and updated according to

$$G_{t+1}(A_t) = G_t(A_t) + \alpha_G G_{t+1}(A_t)\delta_t,$$
$$N_{t+1}(A_t) = N_t(A_t) - \alpha_N N_{t+1}(A_t)\delta_t.$$

for the selected action $A_t$ with $G_{t+1}(a) = G_t(a)$ and $N_{t+1}(a) = N_t(a)$ for actions $a$ not selected and with $G_0 = N_0 = 1$. For simplicity, we will use $\alpha = \alpha_G = \alpha_R = 0.1$ [6].

There are notable differences between the update for $G$ and $N$ and the updates of our model. The first is that the updates of $G$ and $N$ are proportional to $G$ and $N$. This feature was argued to both better reflect Hebbian learning, keep $G$ and $N$ positive, and introduce greater asymmetry into the relationship between $G$ and $A$. Mathematically, however, these updates can be unstable in several ways. First, if $\delta_t > 1/\alpha$ such as would happen if there were large rewards, then $N$ would become negative. In addition, $G$ can grow without bound, even when prediction errors $\delta_t$ are zero on average.

Another key difference is that the OpAL model is relatively insensitive to reward uncertainty compared to our Competing-Critics model. Recall, reward uncertainty plays a central role in our model, since it influences an individual’s sensitivity to both risk and uncertainty when making decisions. To demonstrate this

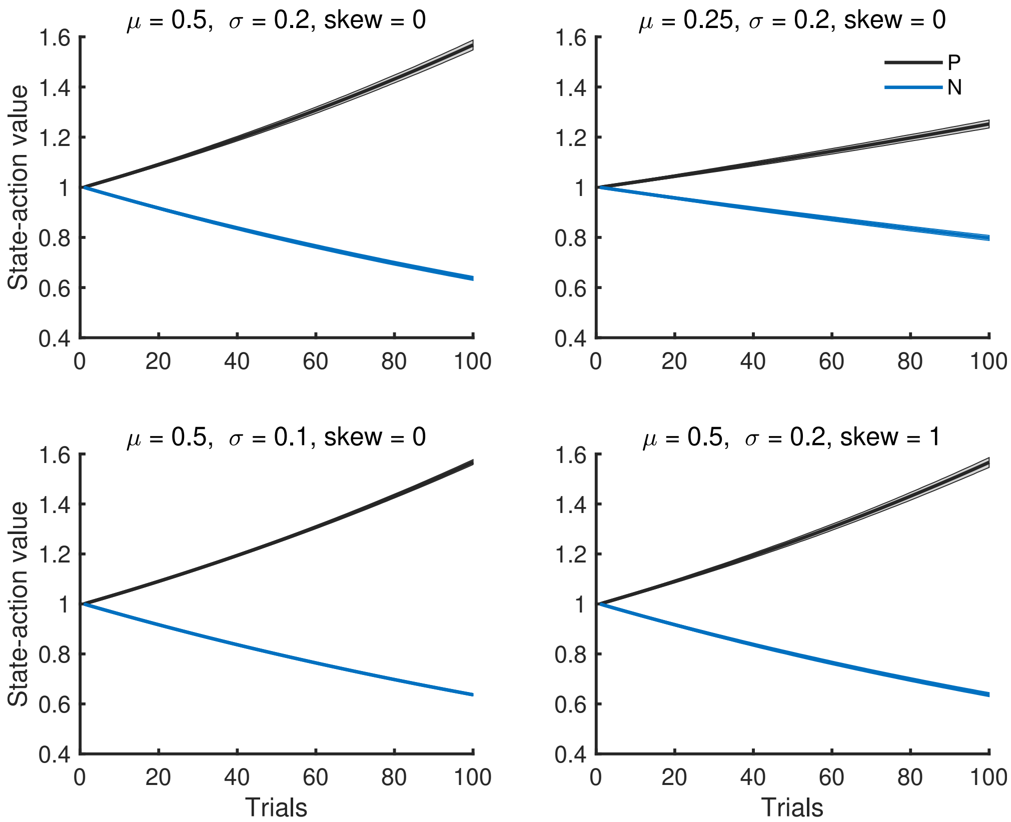


**Fig 1.** Mean and interquartile range for $G$ and $N$ in the OpAL model.

difference, we simulate the OpAL model on our learning example in the main text. First note the exponential growth of $G$ (Figure 1). In addition, note that the curves for $G$ and $N$ only change when the mean is changed, in which case the distance between $G$ and $N$ is cut in half. Thus, $G$ and $N$ are insensitive to the standard deviation $\sigma$ of rewards, whereas $Q^{\pm}$ reflect changes in $\sigma$ in our model.

When using the OpAL model to make decisions, the author propose that decision $a$ is selected with probability proportional to

$$e^{\beta_G P(a) - \beta_N N(a)}$$

for constants $\beta_G$ and $\beta_N$. Thus, despite having two values for each decision, only a weighted difference between these values matter when making a decision. This scalarization is also what is related to reaction time in the OpAL model [6]. By contrast, there is no way to transform $Q^+$ and $Q^-$ into a single value, upon which decisions and reaction times are made in our model.

Further, since $P$ and $N$ are relatively insensitive to reward uncertainty, then decision-making behavior is relatively insensitive to risk. With equal constants $\beta_G = \beta_N = 1$, for example, the OpAL model prefers the

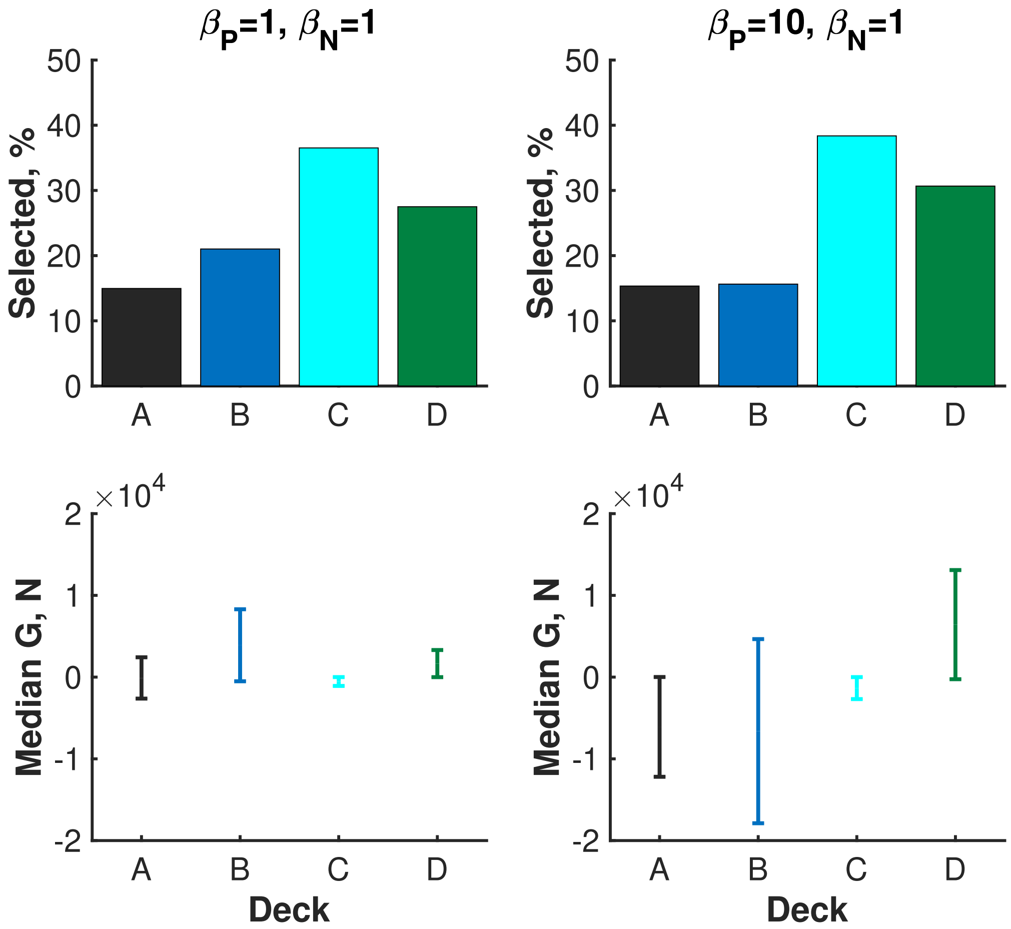


**Fig 2.** Median values of $G$ and $N$ and probability of desk selection during the Iowa Gambling Task in the OpAL model.

"good" deck C in the Iowa Gambling Task rather than the more risky Decks A and B (Figure 2). Increasing the weight $\beta_G$ on the "go" population from 1 to 10 only reinforces the choice of Deck C rather than activating more risky decisions. We also point out the large values of $G$ due to exponential growth and the negativity of $N$ due to the large errors $\delta_t$.

The authors of the OpAL model posit that prediction error $\delta_t$ is still captured by dopamine transients [6], but do not similarly relate serotonin transients to model-derived variable. To investigate whether updates to $G$ or $N$ might reflect serotonin transients (particularly the no-go system $N$), we simulated the stock market task with the OPaL model (Figure 3). Overall (left panels), the updates $G$ and $N$ mirror the updates $\Delta Q^+, -\Delta Q^-$ in our model and hence, the trends of the dopamine and serotonin transients in the experiment by Moran *et al* ( [7]). However, when broken down by reward prediction error (RPE), the two models no longer agree. In particular, the update $\Delta N$ is not largest when switching from a high to low bet during negative RPE or from a low to high bet during positive RPE. Hence, $\Delta N$ does not mirror the trends of

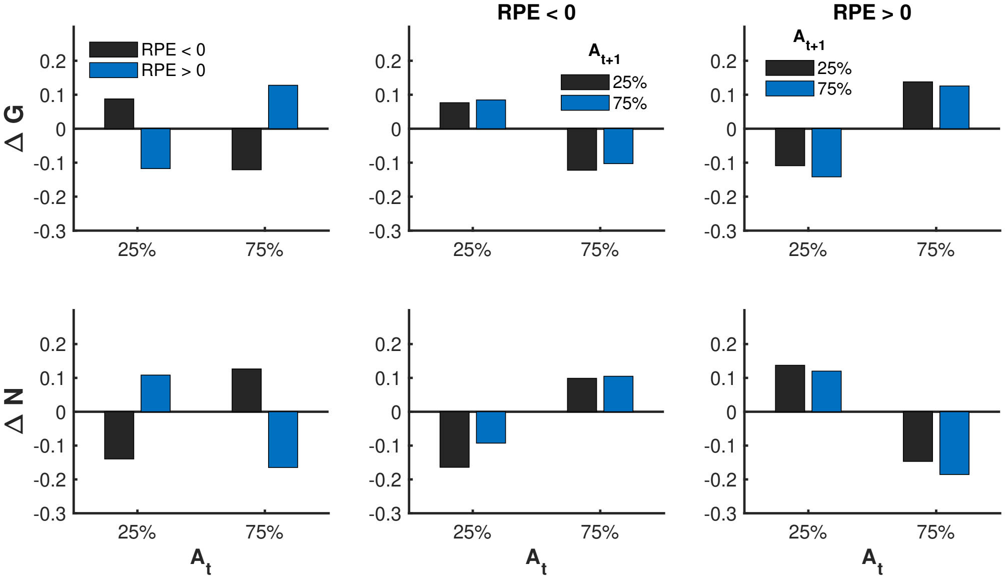


**Fig 3.** Mean updates of $G$ and $N$ during the stock task in the OpAL model.

serotonin transients in [7], where a relatively large serotonin transient preceded a lowering of a bet when RPE was negative and preceded a raising or holding of a bet when RPE was positive.

In summary, both the OpAL model and our Competing-Critics model use dual learning systems that oppose each other and connect these systems to "go" and "no-go" systems. However, there are a number of notable differences between the models in terms of sensitivity to reward uncertainty and risk, stability, scalarization of dual values to influence decisions and reaction time, and direct relationship between updates and serotonin transients. For these reasons, we believe the Competing-Critics model contributes a new conceptual framework for human decision-making.

### References


1. Watkins CJCH, Dayan P. Q-learning. In: Machine Learning; 1992. p. 279–292.

2. Mihatsch O, Neuneier R. Risk-sensitive reinforcement learning. Machine learning. 2002;49(2-3):267–290.

3. Diaconis P, Freedman D. Iterated random functions. SIAM Review. 1999;41:45–76.

4. Duflo M. Random Iterative Models. Springer Berlin Heidelberg; 1997. Available from: `https://doi.org/10.1007%2F978-3-662-12880-0`.

5. Daw ND, Gershman SJ, Seymour B, Dayan P, Dolan RJ. Model-based influences on humans' choices and striatal prediction errors. Neuron. 2011;69(6):1204–1215.

6. Collins AG, Frank MJ. Opponent actor learning (OpAL): Modeling interactive effects of striatal dopamine on reinforcement learning and choice incentive. Psychological review. 2014;121(3):337.
7. Moran RJ, Kishida KT, Lohrenz T, Saez I, Laxton AW, Witcher MR, et al. The protective action encoding of serotonin transients in the human brain. Neuropsychopharmacology. 2018;43(6):1425.
